# Embodiment of a functionally altered virtual arm in adults and children

**DOI:** 10.1101/2025.01.03.631079

**Authors:** Leif Johannsen, Claire Yuke Pi, Simon Thurlbeck, Marco Gillies, Sylvia Xueni Pan, Dorothy Cowie

**Author notes:** Corresponding author: Leif Johannsen RWTH Aachen University Institute of Psychology Jägerstraße 17-19 52066 Aachen Germany.

## Abstract

While the embodiment of non-human body parts is well-established, the precise conditions that facilitate this remain incompletely understood. One critical yet underexplored factor is whether the non-human body part affords greater or lesser functionality than its biological counterpart. This study investigated how the sensorimotor capabilities of a dynamic virtual arm modulate the experience of embodiment. Given that children’s body representations are widely considered to be more flexible than those of adults, they may be especially suited to embodying functionally altered virtual bodies. To test this, both child and adult participants engaged in goal-directed forward reaching movements with a virtual arm to feed animals within an immersive virtual environment. Reaching functionality was systematically manipulated via changes in visual gain, adjusting the arm’s length and functionality from a normal (100%) condition, to be slightly reduced (80%); slightly increased (120%); or markedly increased (400%). Our findings reveal that extreme alterations in reaching functionality (400% visual gain) significantly reduced subjective ratings of limb ownership, an effect most evident in adult participants. Despite these perceptual disruptions, participants across all ages adjusted their reach kinematics in ways that reflected an integration of the virtual arm’s perceived capabilities with the physical limitations of their own bodies. Interestingly, children responded to the altered embodiment with more cautious and less refined movement strategies than adults, suggesting developmental differences in adaptive motor control over non-human bodies. Moreover, across both age groups, exposure to functionally enhanced virtual arms led to increased subjective estimates of reaching affordances, highlighting the influence of altered sensorimotor feedback on perceived action capabilities. Collectively, these results demonstrate that the sense of body ownership, the accuracy of body representations, and the properties of sensorimotor control are closely associated with bodily function. Moreover, while children may exhibit greater tolerance to functional alterations in terms of perceived ownership, they do not show superior motor control. These findings reveal that sensorimotor function and developmental factors interact to shape the boundaries of embodiment in virtual contexts.

## Introduction

Our physical and sensorimotor abilities are a highly skilled, evolved system which allow us to use our body to interact in a highly adapted manner within our environment. Naturally, our bodies are also subject to physical limitations. However, recent technological advances allow users to experience augmented bodies. These may be *functionally* different to one’s own - able to reach further, exert greater force, or even fly through virtual space. This raises several related questions: first, what the outer limits are of what we can accept as our own body? Second, how does body function determine the user’s control of the body and their feeling of ownership over it? An important case of augmentation is virtual bodies, which can be used in experiences focused on education (Bailey & Bailenson, 2017), entertainment, and therapy (Won et al., 2017). In this study we ask how users can learn to embody altered virtual bodies. ‘Embodiment’ is an umbrella term for feelings of ownership and agency over the body; and its incorporation into one’s own motor control schema. We focus on how the functionality of a virtual limb in a specific context may limit its embodiment. We explore this not only in adults, but also in children, whose high levels of neural plasticity and over-reliance on visual signals for locating their own body (Cowie et al., 2013) make them a key group for mapping the outer limits of body augmentation.

Understanding one’s own physical body arises from a complex interaction of multisensory signals and prior expectations. In this intricate system, there is some plasticity in what we can accept as our own body. For example, a spatially displaced, plaster-cast rubber hand which is stimulated at the same time as our own can elicit subjective feelings of ‘ownership’ (Botvinick & Cohen, 1998; Gonzalez-Franco & Peck, 2018), with corresponding neural activity (Ehrsson, 2007) and a drift in perceived own-hand position. Such bodily illusions can be maintained for fake hands which differ in texture (Preston & Kirk, 2022) or size (Cowie et al., 2022; Pavani & Zampini, 2007): therefore, there is plasticity in these aspects of own-body representation. Supernumerary body parts such as an extra thumb (Kieliba et al., 2021) or virtual third arm (Guterstam et al., 2011) can also be integrated into the functional repertoire and experienced as part of one’s own body. Bodily illusions can be especially powerful in children under 10 years, who are not only subject to substantial bodily growth, but are more ‘visually captured’ than adults by fake hands (Allen et al., 2023; Cowie et al., 2018; Cowie et al., 2016; Dewe et al., 2022; Gottwald et al., 2021). Therefore, children under the age of 10 years may have especially high levels of plasticity in own-body representation. In sum, we argue that the sense of self – which lies at the core of our experience, memory, and identity - can be manipulated, including in new digital contexts like virtual reality, and especially in children.

However, it is also clear that there are limits on what we can accept as part of our own body. In comparison with a hand-like form, illusions of ownership are drastically reduced for a wooden block (Tessari et al., 2010), misoriented hand (Costantini & Haggard, 2007), or abstract virtual shape, even in a dynamic context (Ferran Argelaguet et al., 2016; Lenggenhager et al., 2007). Likewise, even after years of using a prosthetic, it may be represented significantly differently to a hand by the visual cortex (Maimon-Mor & Makin, 2020). In broad terms, these limits on plasticity can be explained by supposing that novel body parts which differ greatly from one’s own original configuration make unmanageable demands on cognitive, attentional, and/or neural mechanisms (Makin et al., 2017), resulting in a rejection of the novel body part: either through a series of hierarchical comparisons between the original and novel bodies (Tsakiris, 2010), or - in a more recent conception - through a dynamic set of constraints within a probabilistic framework (Apps & Tsakiris, 2014; Chancel et al., 2022). From this body of work is it apparent that while body representation has a degree of plasticity, it also has limits.

Given this balance of possibility and constraint, and with the increasing availability of both real and virtual augmented bodies, it is important to evaluate where the bounds of augmentation lie. In particular, it seems key to uncover the dimensions on which such limits might operate. Posture is certainly one such dimension: minor discrepancies between felt and seen hand postures can severely disrupt bodily illusions at all ages (Gottwald et al., 2021; Tsakiris & Haggard, 2005). A dimension referred to as ‘form’ or ‘corporeality’ is clearly important, in that embodiment is reduced for a wooden block (Tsakiris et al., 2010) or moving hand with missing fingers (Schwind et al., 2017). Yet current work does not clearly explain why despite these form limits, users can accept a third arm (A. S. Won et al., 2015), or lobster body (Andrea Stevenson Won et al., 2015). In this paper we introduce a theory which may reconcile these seemingly incoherent findings. We propose that a key dimension determining embodiment is body functionality: our ability to embody an object depends on how well it allows us to achieve a functional end or goal. On this view, reduced function which prevents participants from embodying a wooden block or minimal hand; while enhanced function permits embodiment of a third arm or lobster body. While this idea is not without precedent in the literature (Aymerich- Franch & Ganesh, 2016; Cardinali et al., 2021), the role of functionality in embodiment has not been systematically isolated and experimentally tested.

In order to do this, we used a modified ‘GoGo’ technique (Poupyrev et al., 1996) to alter the function of a moving virtual arm. Crucially, we could therefore either reduce or increase its functionality, and indeed we could vary this parametrically. Visual gain (the ratio of virtual to real arm position) was manipulated so that the virtual arm was made to extend far into space or perform shortened reaches. In an ‘increased function’ condition, a reach in real space resulted in an extended reach for the virtual arm: the visual gain between the participant’s reach and the virtual arm reach was greater than one. In a ‘reduced function’ condition, a reach in real space resulted in a shorter reach for the virtual arm: the visual gain between the participant’s reach and the virtual arm reach was below one. We compared baseline experience with several between-subjects conditions: a reduced-function arm with visual gain of 0.8 (‘F-’), an increased-function arm with visual gain of 1.2 (‘F+’), and an increased-function arm with visual gain of 4.0 (‘F++’). For a participant with arm length 60cm, their maximum reach in the F- condition would be only 48cm, while their maximum reach in the F++ condition would be 240cm. To allow a direct comparison of reduced and increased function, the F- and F+ conditions were symmetrical about 1. To compare different degrees of enhanced function and explore the upper limits of plasticity, we compared the F+ and F++ conditions. Finally, we tested not only adults, but also two groups of children: we predicted that younger children (5-7 years), with less experience of their own body, might be better able to feel ownership over the virtual arm than older children (8-10 years) or adults. In contrast, we proposed that skilled control might be easier for the older 8-10-year-old children and adults, who are better able to integrate sensory feedback for motor control (Hay, 1990).

We used multiple indices to test how functionality constrained embodiment of the arm. This allowed us to establish how functionality affected all relevant aspects of body representation, and to reveal differential effects across ownership and the motor schema, which are often independently affected by bodily illusions (Rohde et al., 2011). Subjective experiences of ownership and agency were measured through questionnaire. We hypothesized that relative to a typical body, ownership would be maintained for bodies with slightly altered function and reduced for those with more strongly altered function. This range of embodied forms was expected to be smaller for reduced-function than enhanced-function bodies; and children were expected to show greater subjective embodiment of highly altered bodies than adults. Motor incorporation was measured firstly by pre- and post-experience affordance judgments of how far the users could reach with their own arm (L. P. Lin et al., 2020). As well as being of significant practical importance, such after-effects of the virtual experience would show a deep level of incorporation, which we predicted to be greatest in children.

Motor incorporation was secondly assessed with kinematic indices of visuomotor control. While reach smoothness functioned as a general indicator of good control, the other measured variables centre around the fact that a reach comprises two phases, either side of a velocity peak: an “approach” phase of preplanned, accelerative movement towards a target, followed by a “homing-in” phase of feedback-driven deceleration (Georgopoulos, 1986; Jeannerod, 1988). Skilled motor incorporation of the virtual arm would be shown by a key characteristic of real world reaching, namely an increase in peak velocity with target distance, yielding efficient, faster reaches for further targets (Ojakangas & Ebner, 1991). However, for highly-functionally- altered arms we might expect some dampening of the pattern to account for the limited reach of the physical body.

Further, the spatial and temporal positioning of the peak can be considered a specific reach solution (Jakobson & Goodale, 1991) which reveals the degree and manner of motor incorporation. For example, in a real-world reach, an invariant spatial or temporal proportion of the reach across different target distances is considered an efficient basic solution, as it relies on a constant motor control scheme that simply scales to distance (e.g. always 50% of reach time or distance; (MacKenzie et al., 1987; Schmidt, 2003). In different contexts, peak position might shift, with an earlier peak representing greater emphasis on caution and accuracy, and a later peak greater confidence and speed (Peternel et al., 2017; Plamondon & Alimi, 1997). We predicted that peak positions might remain unchanged for small changes in functionality but reveal some warranted caution in the extreme (F++) case. Likewise, we expected to find more cautious early peaks in children than in motorically skilled adults.

## Results

Participants completed an animal-feeding reaching task within an immersive virtual environment (Fig. 1, Fig. S1). Each participant completed a baseline block of reaches in which the virtual arm extended as the real arm, followed by an experimental ‘GoGo’ block where the arm’s function changed. In the GoGo block, the virtual arm position was a multiple of the real arm position depending on condition (Fig 1A): 0.8 (F-), 1.2 (F+), or 4.0 (F++).

**Figure 1.**
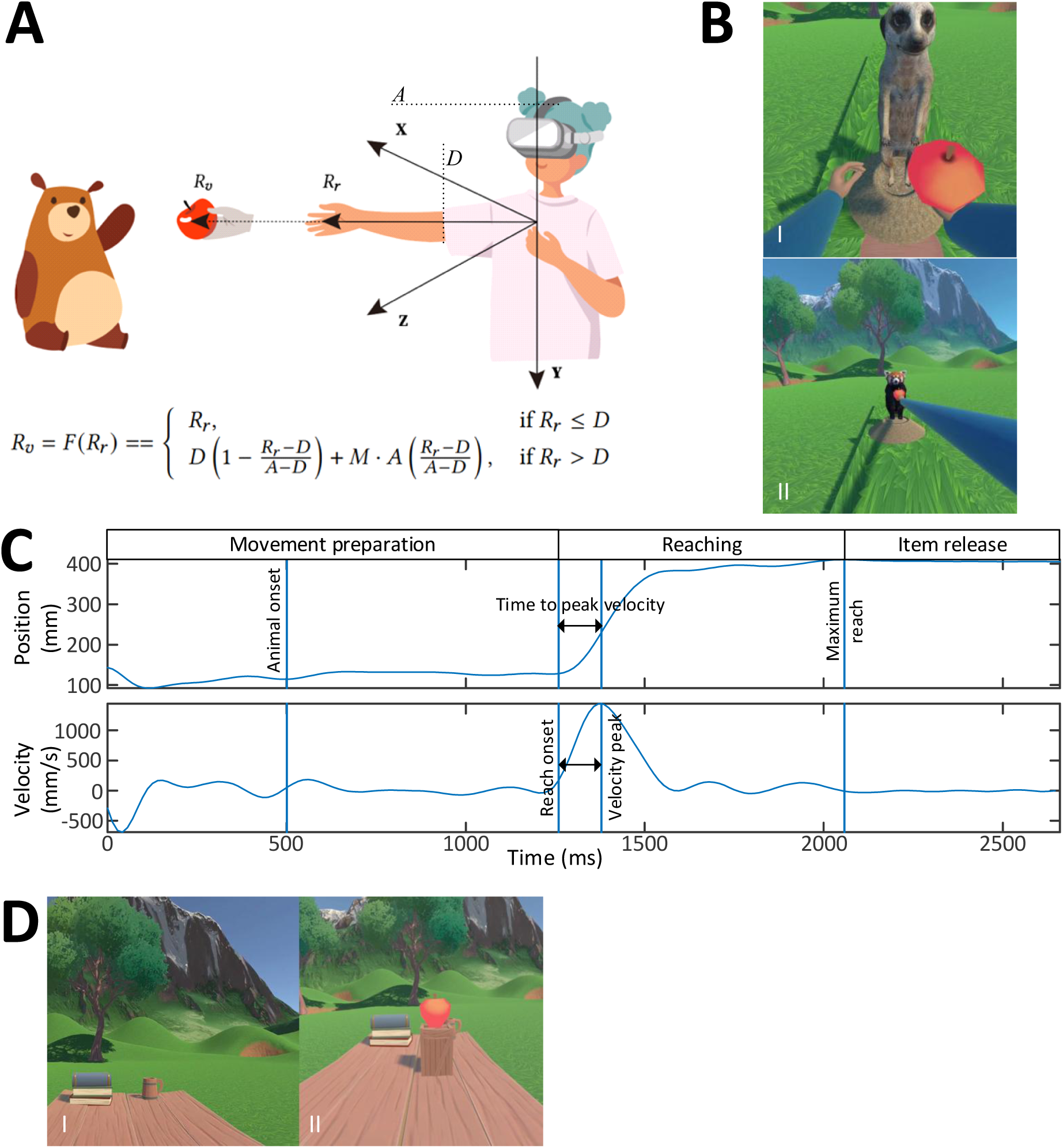
A.) a relationship between real (Rr) and virtual (Rv) hand position: an inflection point (D) at 2/3 arm length (A) was chosen as a comfortable distance for a transition between normal and altered functionality. As the real arm extended further than D (R𝑟 > D), the virtual arm length began to change in linear proportion to the real distance travelled. The maximum was a multiple, M, of the arm length: in the F- condition M=0.8; F+ M=1.2; F++ M=4.0. B.) The reaching sequence for a normal (I) and a F++ (II) reach: moving into a ready volume triggers the appearance of a food item, grasping this triggers onset of a target animal, the forward reach is performed and the food released, the animal eats and food disappears. C.) Anteroposterior position and velocity for an illustrative reaching trial of a single participant. Vertical lines indicate (from left to right) the movement preparation period and onset of the target animal, reach start, peak velocity, time to peak velocity, and reach end D.) Reaching affordance task. An apple on a pedestal is presented; no virtual body is visible. The participant verbally judges if it is or is not reachable with their real arm.

Each of 259 participants experienced only one altered function condition. Age groups were young children (5-8 years, N = 70), older children (8-11 years, N = 101), and adults (N = 88). Reaches were always made to four distances: two before the inflection point, at 52 and 64% of each participant’s fully extended arm length; and two after the inflection point, at 88 and 100%. The 64% arm length target distance was chosen as it was in the normal range just below the inflection point at 66% arm length. Similarly, the 100% arm length target distance was chosen at the very end of the GoGo range beyond the inflection point. The remaining two target distances were 12% arm length shorter than these values (52% and 88%). The 52% target was deemed suitable as it was about halfway between the start and the inflection point. The 88% target ensured that the closer boundary of the target volume, where the fruit item could be released, was still beyond the inflection point. The actual reaching movements required to reach targets were therefore constant across baseline and GoGo blocks. However, the same reach would appear further in virtual space for F+ and F++ conditions and nearer for F-. Experimental factors were therefore Age (young children, older children, adults), Functionality (F-, F+, F++), Block (baseline, GoGo), and Distance (52, 64, 88,100% arm length). During all reaches, movement was measured using the HMD’s inbuilt tracking mechanism, from which we extracted kinematic variables of interest (Borzelli et al., 2025). After each block, reaching affordance was measured in a new virtual environment in which, without reaching, they made a Y/N verbal judgment on whether they could reach an apple on a table in front of them. Using repeated trials in a psychophysical staircase procedure where the apple’s distance was varied, we measured the threshold at which they judged that they could just reach the apple. Finally, participants answered questions on subjective experience, comprised of perceived ownership and agency over the virtual arm, the extent to which it felt tool-like, their user experience, and a control statement to assess suggestibility. Mixed ANOVAs with factors Age, Functionality and Block were used to analyse each subjective experience measure and reaching affordance; while Mixed ANOVAs with factors Age, Functionality, Block and Distance were used to analyse each kinematic parameter (Table 2).

## Subjective embodiment

We highlight key findings for each of the five Likert scale questionnaire items (Figure 2). These test our hypotheses that compared with typical bodies, ownership would be lower for altered-function bodies, particularly reduced-function bodies, and particularly for adults. First, we note that the functionality of the virtual arm affected the participant’s feeling of ownership over it. This is shown in a significant interaction between functionality group and block, F(2,500)=4.46, p=0.01, pη2=0.02, where only the F++ group reported reduced ownership after the GoGo block (t(500)=2.7, p= 0.007; Fig. 2d; other groups both t(500)< 1.26, p>0.21). Therefore, while ownership is possible for arms with either mildly reduced or mildly enhanced function, it is compromised by highly increased function. This partially supports our hypothesis that differently functional bodies would be less well embodied but does not show the predicted asymmetry. Second, age affected ownership of the GoGo arm. To follow a significant interaction between age group and block, F(2,500)=4.03,p=0.02, pη2=0.02, we examined with Post-hoc comparisons the effect of the GoGo experience on each age group. GoGo reduced ownership for adults (t(500)=3.83, p=0.0001) but not for children (both groups t(500)<1.42, p> 0.16), Fig. 2c). Further, an age by functionality interaction, F(4,500)=4.09, p=0.003, pη^2^=0.03, revealed that adults, but not children, gave lower ratings in the F++ group than in the F- (t(500)=4.34, p<.001) and F+ (t(500)=2.83, p=0.005) groups. These findings partially support our hypothesis that the limiting effects of function on ownership were most pronounced for adults, and less pronounced for children.

**Figure 2.**
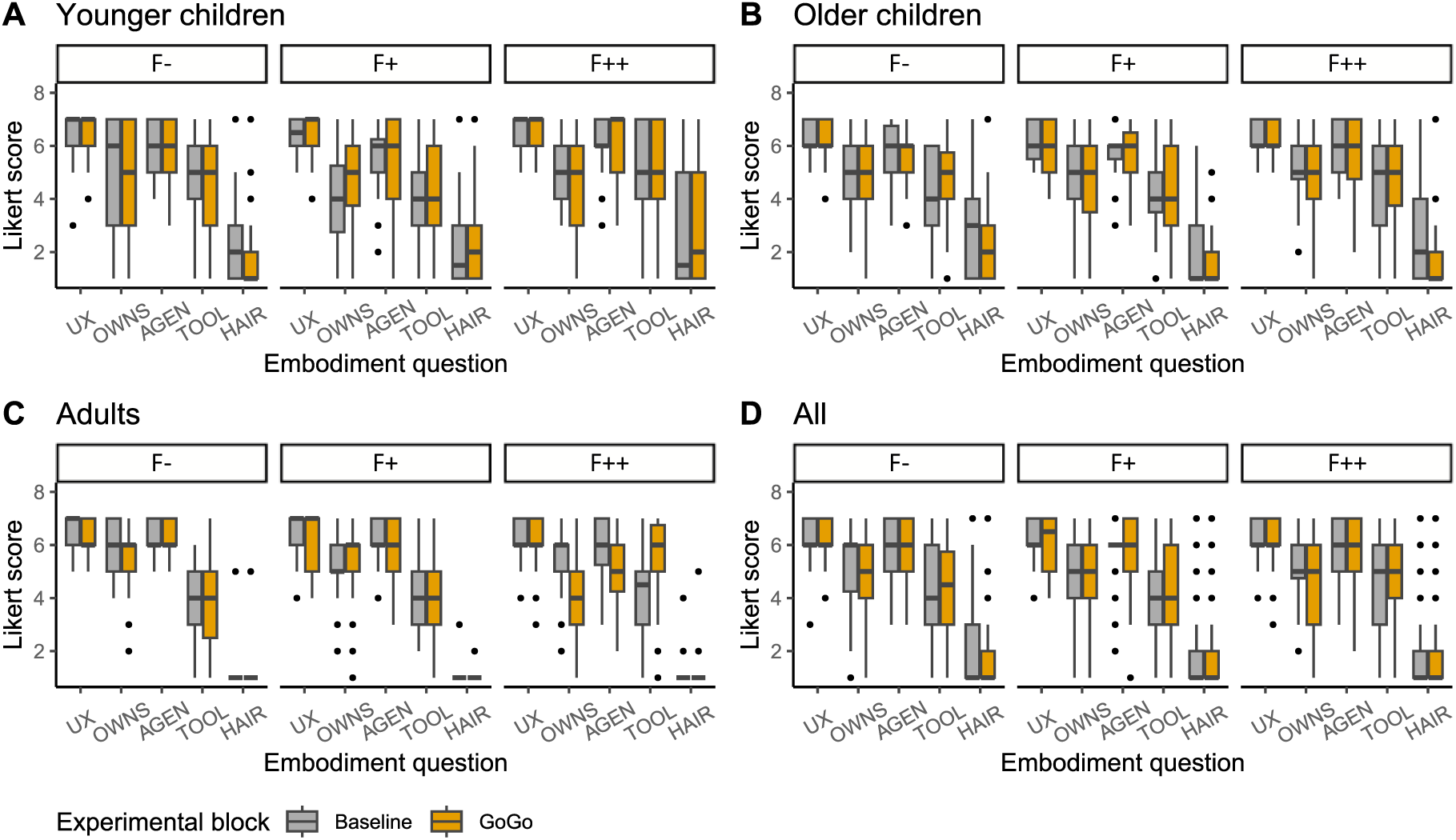
Subjective embodiment ratings as a function of age group, functionality group and block: A) younger children, B) older children, C) adults, D) all three groups. F-: reduced function; F+: increased function; F++: strongly increased function. UX: user experience; OWNS: sense of ownership; AGEN: sense of agency: TOOL: sense of tool-likeness; HAIR: suggestibility control.

While ownership was reduced only in the F++ group, the sense of agency showed a main effect of block, F(1,500)= 8.67,p=0.003, pη^2^=0.02, being slightly reduced across all GoGo conditions (mean=5.60, SD 1.27; t(500)=2.94, p=0.003) compared to baseline (mean=5.95, SD 0.98). (An additional interaction involving age independent of block (F(4,500)=3.59, p=0.007, pη^2^= 0.03) is reported in supplementary materials.) Suggestibility ratings - agreement that a participant’s hair turned blue - were overall low. As expected based on previous findings (Cowie et al., 2018), they were also lower in adults (mean=1.13, SD 0.60) than in children (younger: mean=2.50, SD 2.0; older mean=2.14, SD 1.49; both t(500) >10.34, both p <.0001; Fig. 2; F(2, 500)= 90.13, p<0.001, pη^2^= 0.26). An unexpected 3-way interaction, F(4,500)=2.51, p=0.04, pη^2^= 0.02, was caused by older children in the F++ group reducing their ratings further compared to baseline (t(500)=2.05, p=0.04).

Several additional measures were taken to help contextualise the experience. Function affected tool-likeness ratings, F(2,500)=5.70, p=0.004, pη^2^=0.02: participants in the F++ groups rated the virtual limbs more as resembling a tool (mean=4.79, SD 1.69) than the F- (mean=4.26, SD 1.69; t(500)= 2.88, p=0.004) or F+ (mean=4.32, SD 1.64; t(500)=2.97, p=0.003) groups. User experience was affected by age, F(2,500)=4.82, p=0.008, pη^2^=0.02, younger children (mean= 6.35, SD 0.89) and adults (mean= 6.24, SD 0.89) rated the game slightly more favourably than older children (mean= 6.12, SD 0.78), who were most familiar with videogame technology (mean usage for younger children: 7.2 h/wk, SD 7.2; older children: 9.4 h/wk, SD 8.0; adults: 4.8 h/wk, SD 8.2).

## Motor control

### Peak velocity

In a well-calibrated reaching movement, peak velocity will calibrate with target distance. Across blocks, that was true for our participants’ reaches (52% target distance: mean= 420 mm/s, 64% TD: mean= 488 mm/s, 88% TD: mean= 591 mm/s, 100% TD: mean= 625 mm/s). The effects of target distance on peak velocity also grew stronger with age (age by distance interaction, Table 1): this is explored further in analysis below.

**Table 1.** Overview of significant effects and interactions for each kinematic parameter. Age: Age group; Funct.: Functionality; Block: Experimental block; Dist.: Target distance.

|  | Parameter |  |  |  |
| --- | --- | --- | --- | --- |
|  | Peak velocity | Relative time to peak velocity | Relative distance to peak velocity | Smoothness |
| Age | **F(2,250)=17.17,**<br>**p<0.001,**<br>**$\eta^2=0.12$** | **F(2,250)=7.29,**<br>**p<0.001,**<br>**$\eta^2=0.06$** | **F(2,250)=45.36,**<br>**p<0.001,**<br>**$\eta^2=0.27$** | **F(2,250)=66.98,**<br>**p<0.001,**<br>**$\eta^2=0.35$** |
| Funct. | F(2,250)=2.40,<br>p=0.09,<br>$\eta^2=0.02$ | F(2,250)=1.79,<br>p=0.17,<br>$\eta^2=0.01$ | F(2,250)=1.92,<br>p=0.15,<br>$\eta^2=0.02$ | **F(2,250)=3.92,**<br>**p=0.02,**<br>**$\eta^2=0.03$** |
| Age * Funct | F(4,250)=0.44,<br>p=0.78,<br>$\eta^2=0.007$ | F(4,250)=1.73,<br>p=0.14,<br>$\eta^2=0.03$ | F(4,250)=0.26,<br>p=0.90,<br>$\eta^2=0.004$ | F(4,250)=1.09,<br>p=0.36,<br>$\eta^2=0.02$ |
| Block | **F(1,250)=6.56,**<br>**p=0.01,**<br>**$\eta^2=0.03$** | F(1,250)=0.16,<br>p=0.69,<br>$\eta^2=0.0006$ | F(1,250)=1.07,<br>p=0.30,<br>$\eta^2=0.004$ | **F(1,250)=9.15,**<br>**p=0.003,**<br>**$\eta^2=0.04$** |
| Age * Block | **F(2,250)=6.26,**<br>**p=0.002,**<br>**$\eta^2=0.05$** | F(2,250)=0.02,<br>p=0.98,<br>$\eta^2=0.0001$ | F(2,250)=0.36,<br>p=0.70,<br>$\eta^2=0.002$ | F(2,250)=1.82,<br>p=0.16,<br>$\eta^2=0.01$ |
| Funct. * Block | F(2,250)=0.15,<br>p=0.86,<br>$\eta^2=0.001$ | **F(2,250)=4.58,**<br>**p=0.01,**<br>**$\eta^2=0.04$** | F(2,250)=1.68,<br>p=0.19,<br>$\eta^2=0.01$ | **F(2,250)=19.50,**<br>**p<0.001,**<br>**$\eta^2=0.13$** |
| Age * Funct.<br>* Block | F(4,250)=1.80,<br>p=0.13,<br>$\eta^2=0.03$ | **F(4,250)=3.14,**<br>**p=0.02,**<br>**$\eta^2=0.05$** | **F(4,250)=2.48,**<br>**p=0.04,**<br>**$\eta^2=0.04$** | **F(4,250)=2.87,**<br>**p=0.03,**<br>**$\eta^2=0.04$** |
| Dist. | **F(3,750)=196.76,**<br>**p<0.001,**<br>**$\eta^2=0.44$** | **F(3,750)=7.61,**<br>**p<0.001,**<br>**$\eta^2=0.03$** | **F(3,750)=107.08,**<br>**p<0.001,**<br>**$\eta^2=0.30$** | **F(3,750)=13.99,**<br>**p<0.001,**<br>**$\eta^2=0.05$** |
| Age * Dist. | **F(6,750)=15.06,**<br>**p<0.001,**<br>**$\eta^2=0.11$** | **F(6,750)=4.29,**<br>**p<0.001,**<br>**$\eta^2=0.03$** | **F(6,750)=13.67,**<br>**p<0.001,**<br>**$\eta^2=0.10$** | F(6,750)=0.23,<br>p=0.97,<br>$\eta^2=0.001$ |
| Funct. * Dist. | **F(6,750)=7.30,**<br>**p<0.001,**<br>**$\eta^2=0.06$** | F(6,750)=0.94,<br>p=0.46,<br>$\eta^2=0.007$ | F(6,750)=0.92,<br>p=0.48,<br>$\eta^2=0.007$ | **F(6,750)=4.07,**<br>**p<0.001,**<br>**$\eta^2=0.03$** |
| Age * Funct. *<br>Dist. | F(12,750)=1.24,<br>p=0.25,<br>$\eta^2=0.02$ | F(12,750)=1.31,<br>p=0.21,<br>$\eta^2=0.02$ | F(12,750)=0.98,<br>p=0.46,<br>$\eta^2=0.02$ | F(12,750)=0.93,<br>p=0.52,<br>$\eta^2=0.01$ |
| Block * Dist. | **F(3,750)=6.69,**<br>**p<0.001,**<br>**$\eta^2=0.03$** | **F(3,750)=3.11,**<br>**p=0.03,**<br>**$\eta^2=0.01$** | F(3,750)=0.87,<br>p=0.45,<br>$\eta^2=0.003$ | **F(3,750)=8.51,**<br>**p<0.001,**<br>**$\eta^2=0.03$** |
| Age * Block<br>* Dist. | F(6,750)=0.17,<br>p=0.98,<br>$\eta^2=0.001$ | F(6,750)=0.42,<br>p=0.87,<br>$\eta^2=0.003$ | F(6,750)=1.49,<br>p=0.18,<br>$\eta^2=0.01$ | F(6,750)=1.98,<br>p=0.07,<br>$\eta^2=0.02$ |
| Funct. * Block<br>* Dist. | **F(6,750)=13.51,**<br>**p<0.001,**<br>**$\eta^2=0.10$** | F(6,750)=0.36,<br>p=0.91,<br>$\eta^2=0.002$ | F(6,750)=1.68,<br>p=0.12,<br>$\eta^2=0.01$ | **F(6,750)=4.47,**<br>**p<0.001,**<br>**$\eta^2=0.03$** |
| Age * Funct.<br>* Block * Dist. | F(12,750)=1.66,<br>p=0.07,<br>$\eta^2=0.03$ | F(12,750)=1.61,<br>p=0.09,<br>$\eta^2=0.03$ | **F(12,750)=2.71,**<br>**p=0.001,**<br>**$\eta^2=0.04$** | F(12,750)=1.40,<br>p=0.16,<br>$\eta^2=0.02$ |

The GoGo experience elicited new motor solutions. There was an overall increase in peak velocity for older children (age by block interaction, baseline: emmean=523.0, SD 141.70; GoGo: emmean=566.0, SD 154.77; t(250)=3.23, p=0.02; Fig 3b). Across groups, GoGo changed the scaling of velocity to target distance (Fig. 3d; Table 1). A 3-way Function x Block x Distance interaction showed that in the GoGo block, the calibration of peak velocity to target distance was altered in the F- group (F(3,249)=8.99, p<0.001, pη^2^= 0.10), and F++ groups (F(3,252)=18.18, p<0.001, pη^2^= 0.18; full statistics in supplementary materials; asterisks indicate significant differences in Fig. 3).

**Figure 3.**
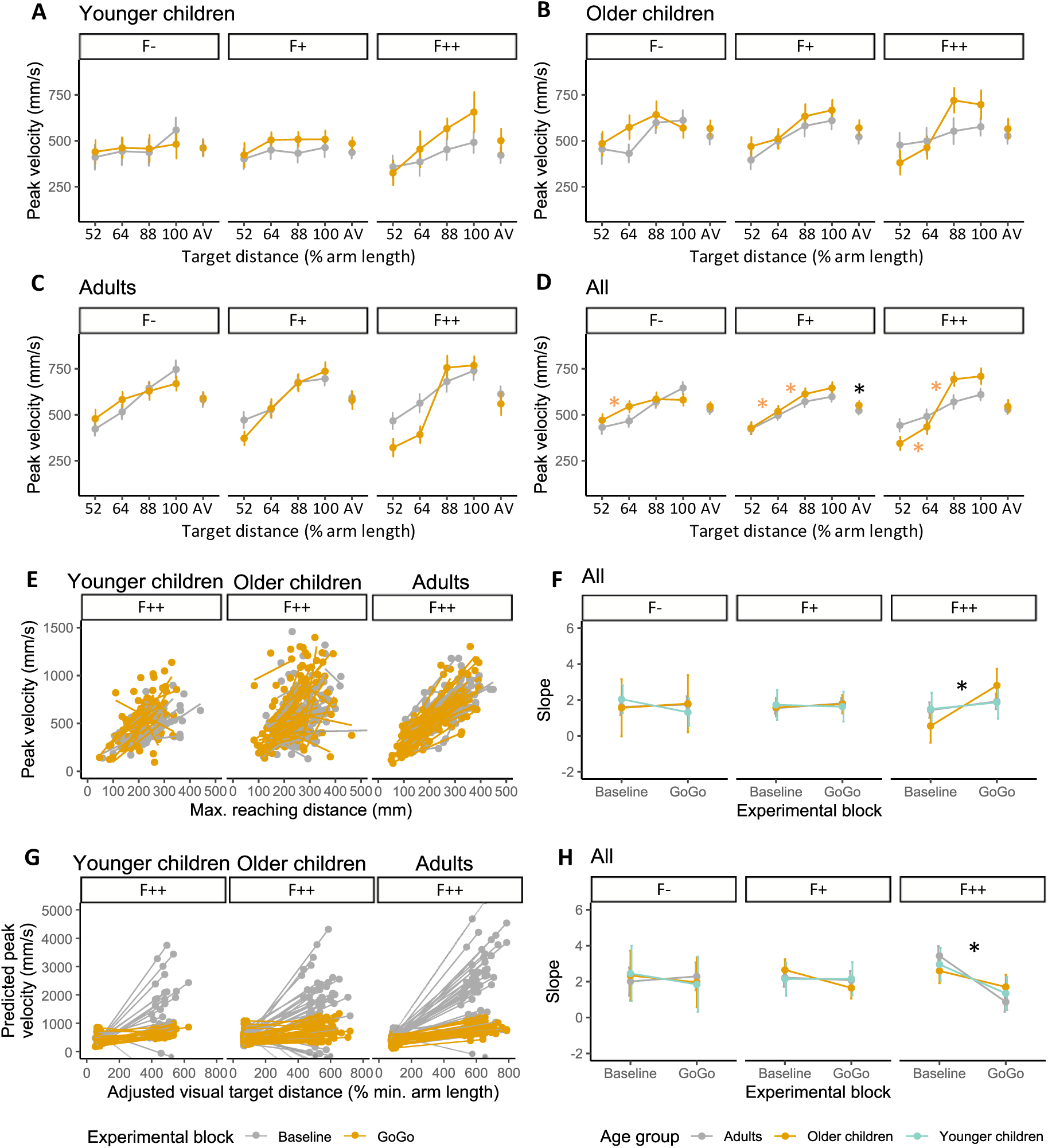
(A – D) Peak velocity as a function of age group, functionality group, block and target distance: A) younger children, B) older children, C) adults, D) all three groups. (E) Linear relationship between peak velocity and maximum reaching distance by block, age and condition (F) expressed as slopes, collapsed across groups. (G) Linear relationship between peak velocity and normalized perceived visual target distance (H) expressed as slopes, collapsed across groups. Panels E and G contrast the predictions of the linear regression model for observed baseline (grey) and GoGo block (orange) velocities. In G, for the F++ GoGo condition, peak velocities do not approach those predicted by the baseline relationship. Error bars show the confidence interval. Asterisks indicate significant single comparisons (p < 0.05). For post-hoc comparisons of interactions, asterisks between neighbouring target distances indicate a significant difference (black: irrespective of block; orange: GoGo only).

To explore these findings, we assumed that participants followed a unitary visuomotor control scheme across the four target distances (a fixed relationship between the distance of a planned reach and its peak velocity); and tested whether this might hold across ages and conditions. To do this we accounted for the fact that adults made physically longer reaches because of their longer arms, performing for each participant linear regression analyses of peak velocity *as a function of reach distance*, where equal regression slopes would indicate the same control scheme. We then conducted ANOVA on these slopes to examine how they changed with Age, Functionality and Block. There was no age effect, suggesting that adults and children indeed adopted similar control schemes. (As reach distance was defined as the maximum reached distance in a trial, it suggests that the distance by age interaction reported in the ANOVA above resulted partly from adults with longer arms reaching further distances and peak velocities.) However, independent of age (Fig. 3f), these slopes showed an interaction between functionality group and block (F(2,250)=3.18, p=0.04, pη^2^= 0.02): in the F++ groups only, the slope was greater in the GoGo block than at baseline (t(250)=3.01, p=0.03; Fig. 3f). Across ages, therefore, we suggest that participants applied a unitary control scheme across baseline, F-, and F+. This suggests that the differences of slope between baseline and GoGo F- reported in the ANOVA above do not hold in this more granular analysis. Most notably, it shows that in the highly functionally enhanced F++ condition participants employed a different scheme with amplified peak velocity.

We next noted that the since the *visual distance* at which the target animals were presented was scaled to arm length, adults reached for visually further targets. This is therefore a potential confounding factor regarding age differences. We therefore performed a set of linear regressions of peak velocity *as a function of the visual distance of targets* (supplementary materials). If peak velocity calibration as a function of the perceived visual distance was maintained across ages and conditions, the regression slopes would be equal. However, they were different between the age groups (F(2,250)=10.52, p<0.001, pη^2^=0.08) with adults (emmean=3.04, SD 2.54) showing steeper slopes than older children (emmean=2.08, SD 2.54; t(250)=2.60, p=0.03) and younger children (emmean=1.18, SD 2.54; t(250)=4.57, p<0.001). Adults therefore increased peak velocity more for *visually far* targets than the children. This general property of their reaches in the virtual environment does not reflect their specific adaptation to altered-function arms.

Focusing specifically on effects of function, the regression slopes on visual distance showed effects of functionality group (F(2,250)=5.12, p<0.001, pη^2^=0.04) and block (F(1,250)=12.97, p<0.001, pη^2^=0.05) as well as an interaction between these factors (F(2,250)=4.72, p=0.01, p<0.001, pη^2^=0.04). Specifically, the GoGo block in the F++ condition had a lower slope (emmean=0.65, SD 3.36) than baseline (emmean=2.34, SD 2.83; t(250)=4.55, p<0.001; Fig. 3h). Therefore, while in the F++ condition participants adopted higher peak velocities as reaching distance increased (Fig 3f) because they calibrated reaches based on visual target distance, they did not ramp up velocity in an unrestricted fashion, achieving extremely high speeds for 400% arm length virtual targets in the F++ condition. Rather, they allowed proprioception of the real arm to moderate their movements even as they responded to the visual pull of the virtual targets and extended arm.

### Relative time to peak velocity (rTPV)

Recall that the velocity peak divides the reach into an accelerative, pre-planned phase before the peak, followed by a decelerative control phase. A short temporal proportion of the reach before peak velocity (rTPV) therefore shows a cautious, longer time left for decelerating control. Across target distances, an efficient basic control scheme holds this peak position constant in relative terms. Indeed, ANOVAs to resolve the interaction between age and target distance showed that younger children (F(3,201)=1.31, p=0.27, pη^2^=0.02; BF=0.04) held rTPV constant. In contrast, older children (F(3,294)=4.02, p=0.008, pη^2^=0.04; BF=0.66) and adults (F(3,255)=19.45, p<0.001, pη^2^=0.19; BF>100), reduced rTPV with increasing target distance (Fig. 4d), leaving proportionally longer control phases for far targets. This age-dependent reaching pattern in the virtual environment was maintained across baseline and Gogo blocks. (In fact - see supplementary materials - adults and by tendency older children held *absolute* timing constant, which resulted in the observed reductions in *relative* timing.)

**Figure 4.**
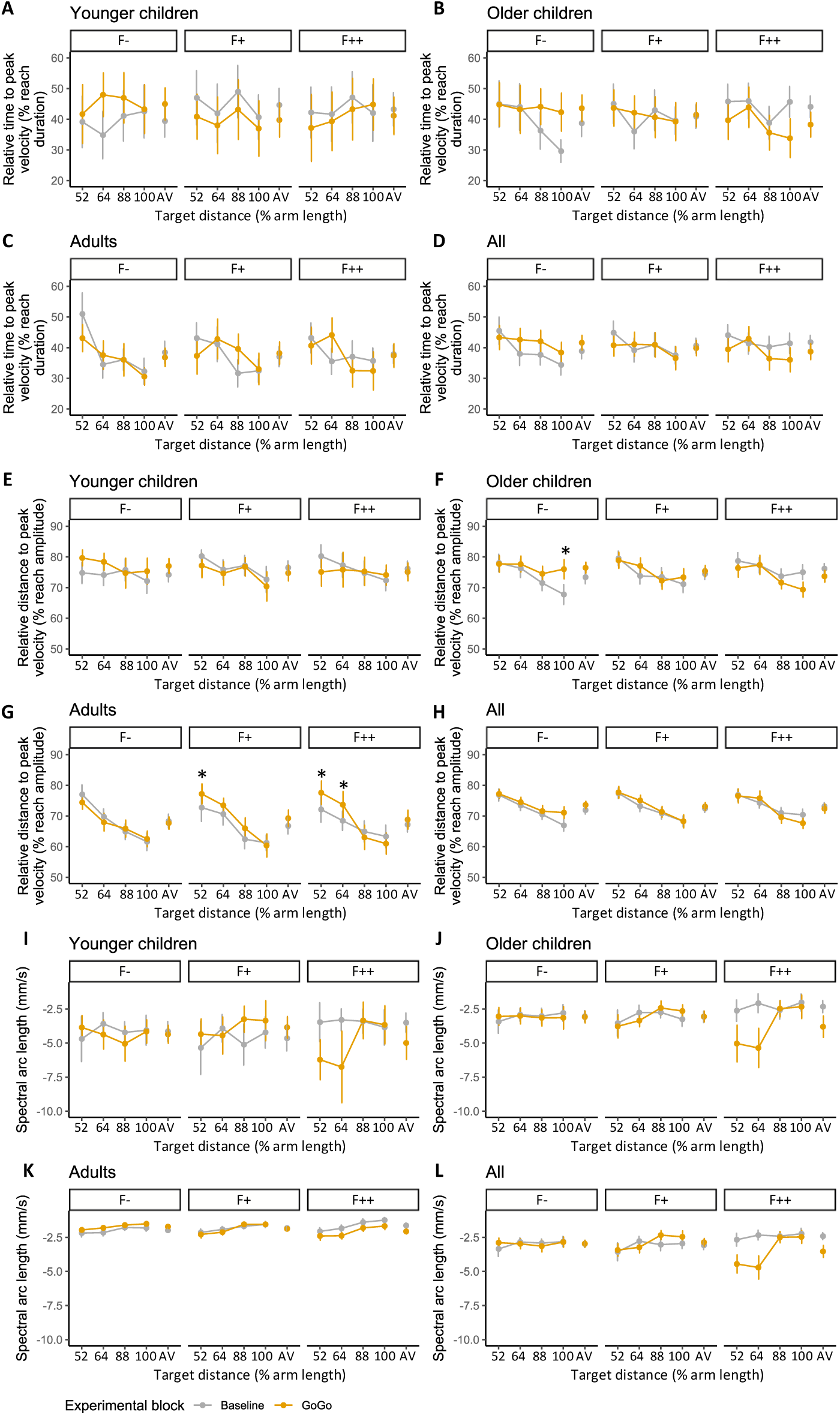
(A-D) Relative time to peak velocity as a function of age group, functionality group, block, and target distance: A) younger children, B) older children, C) adults, D) all three groups. (E-H) Relative distance to peak velocity as a function of these factors. (I-L) Arm extension smoothness as a function of age group, visual gain group and block. Error bars show the confidence interval. For post-hoc comparisons of interactions, asterisks between neighbouring target distances indicate a significant difference (p < 0.05). Asterisks above a specific target distance indicate a main effect of block.

Leaving aside target distance, rTPV showed an interaction between age group, functionality group and block (Table 1; Fig. 4a-c). Follow-up ANOVAs for each age group showed that the parameter was adjusted more to altered functionality by children (change from baseline in younger children: F- = 5.6 %, SD 15.9; F+ = -4.9 %, SD 16.7; F++ = -2.1 %, SD 12.2; change from baseline in older children: F- = 4.9 %, SD 12.3, F+ = 0.5 %, SD 12.6; F++ = -5.8 %, SD 13.7) than by adults (change from baseline: F- = -1.6 %, SD 9.9; F+ = 1.1 %, SD 8.7; F++ = -0.4 %, SD 10.2). Specifically, children in particular adopted a longer acceleration period when functionality was reduced and a more cautious, shorter acceleration period when it was increased while adults retained later peaks even with changes in functionality.

### Relative distance to peak velocity (rDPV)

This parameter again indicates the way in which participants divided the reach into an accelerative and a decelerative phase, but here in terms of *spatial* proportions. Across groups, the peak position (rDPV) reduced for distant targets (Table 1, Fig 4e-g). Unlike rTPV, this did not occur because participants held the *absolute* position constant (Supplementary materials). Rather, participants adopted a ‘mixed’ control solution, peaking somewhat further forward in absolute terms, but not so far as to keep the proportion constant across all target distances.

This pattern was stronger in adults than in children. Adults demonstrated clear decreases in rDPV across target distances, which became prominent with increased functionality. For children - especially younger children – this parameter was less sensitive to altered functionality. Statistically, this is revealed as the four-way interaction between all factors (table 2). Follow-up analyses (Supplementary materials) indicated that while younger children showed a reduced rDPV for increasing target distances (F(3,201)=8.10,p<0.001, pη^2^= 0.11; Fig. 4e), both older groups showed an interaction between functionality group, block and target distance (Older children: F(6,294)= 2.90, p=0.009, pη^2^=0.06; Adults: F(6,255)= 3.90,p<0.001, pη^2^=0.08). These trends were stronger in all adult groups (Fig. 4g), we suggest due to a tendency to adopt GoGo-like later peaks even for near targets.

### Smoothness

The smoothness of a reaching movement depends on the number of subsequent accelerations and decelerations and an ideal reach involves only a single velocity peak only. By comparison, smoothness of a reaching movement is reduced when it contains multiple (local) velocity peaks and especially when it involves one or several changes in movement direction. A reach with multiple changes in direction could be termed erratic or it might reflect the frequent occurrence of spatiotemporal adjustments of a reach towards a target, for example potential target over- and undershooting. The greater the number of velocity peaks in a reach, the longer the length of the enveloping arc of the velocity power spectrum, as a representation of smoothness, becomes (spectral arc length; Balasubramanian et al., 2012). Smoothness showed an interaction between age group, functionality group and block (Fig. 4i-k). Separate ANOVAs showed in each age group an interaction between functionality group and block (younger children: F(2,67)=6.91, p=0.002, pη^2^=0.17; older children: F(2,98)=8.70, p<0.001, pη^2^= 0.15; adults: F(2,85)= 17.90, p<0.001, pη^2^=0.30), resulting from reduced smoothness for the F++ group in the GoGo block at all ages (younger children: t(67)=3.31, p=0.02; older children: t(98)=5.22, p<0.001; adults: t(85)=5.20, p<0.001). In addition, adults in the F- group demonstrated a small increase in smoothness in the GoGo block (t(85)=3.24, p=0.02). Overall, therefore, smoothness was preserved for mildly altered function but compromised by highly altered function.

### Reaching affordance

The distance at which a target was perceived as just reachable depended on age group, F(2,249)=27.08, p<0.001, pη2=0.18, and block, F(1,249)= 27.75, p<0.001, pη2=0.10. Overall, children perceived reaching affordance as longer (younger children: emmean=2,19, SD 0.39; older children: emmean=2.12, SD 0.39) than adults (emmean=1.78, SD 0.39). It was also judged as longer after the GoGo block (baseline: emmean=1.97, SD 0.44; GoGo: emmean=2.09, SD 0.42). Finally, an interaction between functionality group and block, F(2,249)=6.42, p=0.002, pη2=0.05, showed that, independent of age, exposure to arms with increased function increased the reaching affordance (F+: t(249)=3.50, p=0.007; F++: t(249)=5.27, p<0.001; Fig. 5d) while F- arms did not change it.

**Figure 5.**
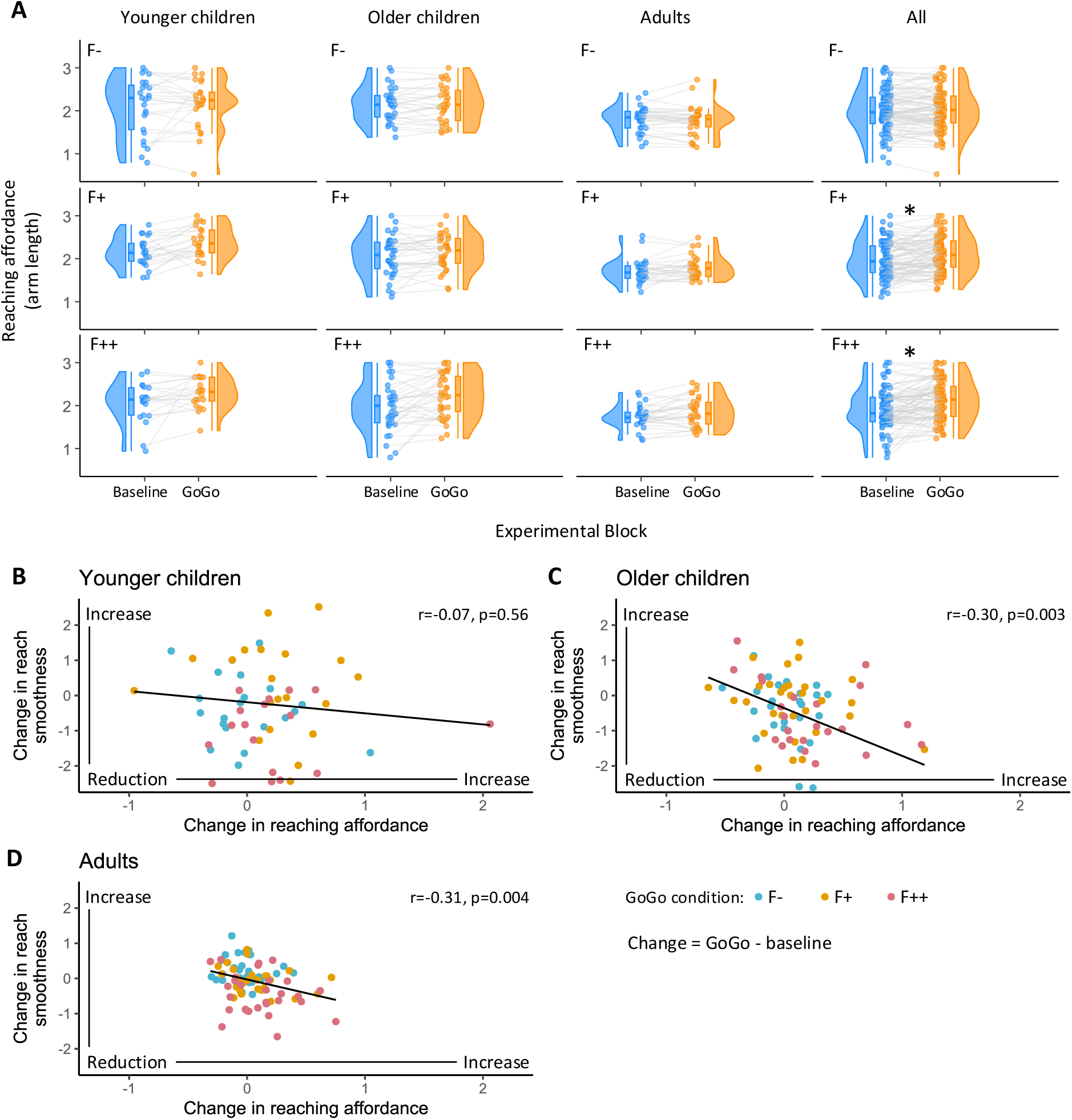
A) Raincloud plots of the reaching affordance as a function of age group, functionality group, and experimental block. F-: reduced function; F+: increased function; F++: strongly increased function. Asterisks indicate statistically significant single comparisons (p < 0.05). Bottom plots show Spearman correlations between the changes in reach smoothness and reaching affordance from the baseline to the GoGo block: B) younger children, C) older children, D) adults. GoGo functionality groups are colour coded.

### Parameter correlations

As expected, different aspects of embodiment remained largely independent, underscoring the importance of measuring these separately and the complexity of the embodied virtual experience. To assess this, we examined a representative measure from each category above. These were ownership (a key measure of subjective experience), smoothness (the kinematic parameter that might most easily be perceptually accessible by participants), and reaching affordance. For each, we calculated the change values from baseline to GoGo block (GoGo value minus baseline value). We then calculated Spearman correlations between pairs of these variables: (i) reaching affordance and the sense of ownership and (ii) reaching affordance and smoothness (to index control efficiency), (iii) smoothness and sense of ownership. These three correlations were calculated for each of the nine age and functionality subgroups: none were significant. To increase power, we next collapsed across the functionality groups to calculate two correlations for each age group. For adults and older children, increased reaching affordance was associated with reduced smoothness from baseline to GoGo blocks (older children: r=-0.30, p=0.003; adults: r=-0.31, p=0.004; Fig. 10a-c). Any correlations between sense of ownership and either reaching affordance or smoothness were not observed for any age group.

## Discussion

### Summary

In the present study, we tested the importance of body functionality to embodiment, by enhancing or reducing functionality using a virtual arm which reached further or less far than the participant’s own. We expected that highly altered function would change several aspects of embodiment, including the sense of ownership, visuomotor control and reaching affordance. We expected that even with a small degree of functional alteration, increased-function arms would be better embodied than reduced-function arms. Finally, we predicted that for children, the sense of limb ownership would be affected less by altered arm functionality, while adjustments in motor control would entail different sensorimotor solutions to those adopted by adults.

### Embodiment & reaching affordance are constrained by function

In line with our hypotheses, a sense of ownership was maintained for bodies with slightly modified function (F- or F+), suggesting that there is plasticity in the range of body functions which users can embody. Within this range, we found no evidence for differences between reduced (F-) and enhanced (F+) function bodies. However, despite their potential advantages, bodies with extremely modified function (F++) reduced the sense of ownership and increased the experience of the virtual limbs as ‘tool-like’. Past a point, therefore, function did act as a constraining factor on ownership. Further, we found that lower ownership occurred predominantly but not exclusively in the adult F++ group. This implies that a constraining effect of function on ownership is somewhat more pronounced for adults than for children. The sense of agency was meanwhile reduced by subtle changes in either direction to the virtual arm’s function. Therefore, perhaps because of the motoric nature of the functional changes, agency seems more sensitive to function than ownership.

By using a body with clear functional modifications but minimal superficial alterations, the present study therefore provides a direct test of how function affects subjective embodiment. We find that there is a range of altered body functions within which embodiment is maintained; that this range may be slightly narrower for adults than children; and that the range is wider for the sense of ownership than the sense of agency. The observed reduction in ownership between 1.2 (F+) and 4 (F++) arm lengths is broadly in line with both a small-scale augmented reality “GoGo” experience (Feuchtner & Muller, 2017); and a study of a non-extendable long virtual arm (Kilteni et al., 2012). These results motivate reconsideration of previous results in the light of functionality. Low ownership over wooden blocks in desktop bodily illusions (Tsakiris et al., 2010); or abstract hands in VR (F. Argelaguet et al., 2016) may be due to their non-functional nature; while acceptance of virtual tails (Steptoe et al., 2013), virtual third arms (A. S. Won et al., 2015), extra fingers (Hoyet et al., 2016), or even tools (Cardinali et al 2021) may arise because their functionality is high. While these examples also contain active movement, we note that this is not sufficient for a sense of ownership when the form is less functional than a human hand (Lin & Jorg, 2016; Argelaguet, 2016). We therefore suggest that functionality is a key dimension affecting embodiment, and that slightly raised functionality can enable embodiment even when superficial morphological characteristics differ from the normal human body.

As predicted, the Gogo experience did not just change the sense of embodiment, but also the perception of own-body functionality, or body schema, as indexed by reaching affordance. We found that judged reachable distance increased following both the enhanced-functionality conditions (F+ and F++), but not the reduced functionality (F-) condition. This suggests that affordance adjustments may occur following both subtle and extreme *increases* in body functionality, while the status quo is maintained response to *decreased* functionality. The increased reaching affordance we observed with a longer virtual arm is in line with previous studies on long virtual arms (D’Angelo et al., 2018; Le Jeune et al., 2024; L. P. Y. Lin et al., 2020), or tools that functionally extend reaching distance (Cardinali et al., 2009; Ladavas & Serino, 2008). This shows that body schema adjustments occur both in contexts where the length of the virtual limbs is held constant (D’Angelo et al., 2018; Le Jeune et al., 2024) as well as in dynamic contexts like our present study. The unchanged reaching affordance we observed with a shorter virtual arm contrasts with Lin et al. (2020), who found reduced reachable distances with fixed-length 50% short arms. The discrepancy may be due to the smaller arm length reduction in our study, and/or the non-extendable arms used in theirs. Given our own finding that affordance changed in F+ but not F- conditions, we suggest that at least within this range of functions and for this measure, our hypothesis is supported regarding asymmetric plasticity in the body schema, with greater tendency to embody increased-function than reduced-function limbs.

### Motor competence remains high with altered-function arms

Motor performance remained impressively robust throughout the visual-proprioceptive discrepancy introduced in the GoGo blocks. Reaches were fast, well-calibrated to target distance, and relatively smooth, rather than being overly slow or erratic (Balasubramanian et al., 2012; Wong et al., 2021). The temporal synchrony between visual and proprioceptive information therefore seemed sufficient for participants to make the ‘unity assumption’ (Chen & Spence, 2017; Welch & Warren, 1980) that both were caused by the same reaching movement. Performance was not perfect: a smoothness reduction for near targets suggests that the motor control solution used for the enhanced-function range of the reach was sometimes erroneously applied to the whole reach. Further, the association of smoothness with reaching affordance in older children and adults may indicate that uncertainty in sensorimotor control drove a subjective shift in the perceived action boundaries. However, the core finding is that even without in-depth training, participants of all ages can show impressive degree of motor control over functionally altered virtual bodies (see also (Clode et al., 2024).

How did participants achieve this? Our participants calibrated peak velocity to target distance, which is a sign of well-controlled reaching (Flash et al., 2013; Viviani & Schneider, 1991; Yokoyama et al., 2018). Furthermore, once absolute arm length differences were accounted for, this calibration was found to be largely constant from baseline to GoGo conditions. This again indicates good control despite the altered ‘GoGo’ visual feedback. Indeed, since the targets were presented at 52%, 64%, 88%, and 100% arm extension in real physical space, and in each GoGo condition, no motor adjustments were necessary to successfully reach for the targets, and participants successfully retained the same solution across baseline, F- and F+ conditions. In contrast, in the F++ condition, participants of all ages were influenced by the amplified functionality of the virtual arm, executing their reaches with disproportionally faster velocities. Participants therefore found a consistent new motor solution for the functionally altered virtual arm. Examining the relation between *visual target distance* and peak velocity, however, revealed a degree of moderation in this adapted reaching. For the far targets (400% arm length in the F++ condition), participants held back: slopes were shallower than predicted by baseline distance-velocity calibration. Thus, adaptation to the visual virtual arm was carefully offset by proprioception of the actual arm, through well-balanced movements that did not exceed the participant’s actual physical boundaries. Our data therefore show that while experiencing new virtual body functions, participants of all ages maintain baseline solutions where possible; make motor adaptations in extremely altered conditions; and choose safe movement solutions which balance the needs of the physical body with those of the altered virtual body.

Interestingly, the faster reaches we observed in increased-function conditions distinguish our results from observed *reductions* in peak velocity following use of a long tool (Cardinali et al., 2009). In a study with a single target position, velocity reductions were interpreted as the tool inducing perception of a longer arm and relatively closer target position within peripersonal space (Cardinali et al., 2009). However, in examining calibration across a wider space, we observed that the longer GoGo arm in contrast caused increased velocity and reaching affordance at far distances. We suggest that this relationship between perceived arm length and peak velocity should be further investigated.

### Specific motor solutions are age-dependent

In line with previous work (Ishak et al., 2014), there was remarkable consistency across age regarding changes in reaching affordance. Further, the crucial calibration of peak velocity to distance was relatively constant across ages. In contrast, age affected several more nuanced aspects of reach kinematics, revealing that specific motor solutions did vary across ages. A first group of age differences were observed across blocks (regardless of functional alterations to the arm). Adults adopted higher peak velocities - perhaps with an awareness that the animals could be fed by placing the hand within a relatively generous target volume, they chose speed over accuracy. However, they also left longer periods of deceleration after the early velocity peak. Therefore, adults arguably dealt with the speed-accuracy trade-off in more adeptly than children. Adults further scaled up peak velocity more in response to *visual distance* than the children, which could indicate a better integration of visual information for planning a reach within the virtual environment. In terms of how different *target distances* were treated, young children achieved peak velocity at a constant temporal proportion of the reach across all target distances, while adults adapted based on context, achieving peak velocity at a shorter proportional time for far distances (constant absolute time). This simple but efficient motor schema in younger children again contrasts with adults’ more refined, task-dependent solution (Schmidt, 2003; Shea & Wulf, 2005). Overall, therefore, we argue that adults showed more nuanced movement patterns than children, adapting flexibly in demanding task conditions (Golenia et al., 2018); and relying strongly on visual information within the virtual environment. These observations add to the growing literature on reaching within VR for both adults (Clark & Riggs, 2020, 2021; Mangalam et al., 2024) and children (Alrashidi et al., 2024).

A second group of age differences were observed in relation to the GoGo function. Adults reduced the proportional time to peak velocity across target distances at baseline and maintained this solution in the Gogo block. In contrast, older children adopted adult-like scaling at baseline, but during GoGo blocks reverted to the constant TPV control scheme that was adopted across blocks by younger children (who further showed a cautious, early peak for all targets in the increased-function GoGo conditions). A similar pattern was seen in the spatial position of the velocity peak. When confronted with modified functionality, therefore, adults appeared to almost effortlessly maintain control; older children demonstrated a fragile control that adapted less well than adults to new body functions; and younger children started with a simple solution and became yet more cautious with the new virtual arm. We finally note that while smoothness overall increased with age as expected (Kuhtz-Buschbeck et al., 1998; Schneiberg et al., 2002), only adults managed to increase the smoothness of their reaching movements in the reduced-function GoGo condition compared to baseline, thus taking clever advantage of the apparently limiting F- condition. In sum, we find that in terms of reach control solutions, children adapt relatively well to a newly functional virtual arm, but adults maintain nuanced control to an impressive degree. Finally, we note that while children were in general more liable to feel ownership with altered-function bodies, this was not coupled to better control.

### Limitations and future directions

The most apparent methodological limitation of the current study is the non-anatomical rendering of the avatar’s arms, which was not possible within the motion tracking solution. The Likert scale embodiment questions likewise represented a relatively coarse measure; more detailed interviews were not feasible given children’s attention spans. Although children were more suggestible than adults, we note that they did not give uniformly high scores but rather moderated their answers across questions and conditions. A future follow-up study should provide an enhanced embodiment experience across a larger number of trials to determine whether children need longer exposure to the functionality manipulations to generate adult-like reaching solutions; and whether any effects that we observed might be transitory in nature, receding after habituation to the functionality alterations. It would likewise be useful to probe how embodiment changes over a broader range of altered functions. Finally, we note that virtual experiences without significant tactile input cannot be expected to elicit the same kind of embodiment as physical augmentation experiences (Clode et al., 2024; Kieliba et al., 2021).

### Conclusion

Altered-function virtual arms were embodied in terms of subjective feelings of ownership and altered bodily schema (perceived reaching affordance and motor control). This plasticity was, however, limited – indeed, this constraining role of function may help account for previous work on embodiment of nonhuman objects. Compared with adults, children had greater ownership over functionally-altered arms but adopted less subtle motor solutions. We suggest that these findings not only inform VR design but demonstrate the crucial role of function in embodied human experience, and its differential impacts on children and adults.

## Materials and methods

### Participants

For this study, all participants were recruited and tested in the United Kingdom. We recruited 171 children of the age range from 5 to 10 years through a local primary school and public engagement events at the Life Science Centre, Newcastle upon Tyne, United Kingdom, and Durham University. This convenience sample was subdivided into a group of 70 children from 5.00 to 7.99 years (mean age=7.07 yrs, SD 0.78; 42 females) and a group of 101 children from 8.00 to 10.99 years (mean age= 9.24 yrs, SD = 1.39; 44 females). In addition, a group of 88 young adults (students from Durham University recruited for course credit) were included (mean age=22.53 yrs, SD = 4.63; 63 females). Each participant was randomly assigned to one of three subgroups exposed to a specific visual control gain (Younger children: F-=25, F+=24, F++=21; Older children: F-=30, F+=35, F++=36; Adults: F-=31, F+=27, F++=30). Table S1 in the supplementary materials summarizes the demographic information for each of the 9 subgroups. The study was approved by the Durham University Psychology Department Ethics committee (reference number: PSYCH-2023-03-31T14_17_04) and performed in accordance with the standards of the 1964 Helsinki declaration of ethical principles for medical research involving human subjects and its later amendments. Written informed consent was provided by all participating adults and by all parents or legal guardians of child participants.

### Materials and equipment

The virtual reality scene was custom-built in Unity and presented by means of a Meta Quest 2 headset. It was an outdoor scene (Fig. S1c): a meadow encircled by tall peaks, with trees and a flowing brook. A grassy mound was visible directly in front of the participant: target animals appeared from here, at a constant height which made leaning forward unnecessary. Further, participants were instructed not to lean forward or to stand during the virtual experiences. Finally, again to ensure a fixed position, child participants sat in a modified car seat with an oriented backrest and body height-adjusted headrest, mounted on a wooden platform (Fig. S1b). They were strapped in with a pair of diagonal seatbelts. Adult participants sat in a standard chair with armrests and were strapped in using a belt at chest height.

The participant’s virtual avatar kept a seated posture on a stool. Participants could not see its trunk, lower body, or legs. The participant’s virtual hands were rendered based on the headset’s hand tracking and gesture recognition features. Two blue sleeves represented the arms, which connected the hands to the avatar’s shoulders with simplified ‘tubes’. The choice not to articulate the arms in an anatomically correct way (e.g. with wrist and elbow joints) allowed for a simple, portable motion capture solution.

### Experimental design and protocol

Data collection was performed in a single session of ∼20 minutes divided into two experimental blocks. The first block, which was identical for all participants, comprised practice trials and baseline data collection. The second block assessed participants during and after the experience of reaching with a functionally different ‘GoGo’ arm. In this block, one third of participants had a mild functional reduction (*F-*, 0.8 visual gain); one third had a mild functional enhancement (*F+*, 1.2 visual gain); one third had a strong functional enhancement (*F++*, 4.0 visual gain). In contrast to Poupyrev and colleagues (1996), we chose a linear relationship between real (Rr) and virtual (Rv) hand position (see equation, Fig. S1a), because a linear relationship allowed the maintenance of a constant scaled gain of maximum arm length. Based on piloting, an inflection point (D) at 2/3 arm length was chosen as a comfortable distance for a transition between normal and altered functionality. When the user extended the arm further than D (R𝑟 > D), the virtual arm length began to change in linear proportion with the real distance travelled.

Before the start of the reaching task, we took participant’s age and average video game experience; and measured their body height and forward extended arm length manually, as well as electronically by the virtual reality headset. Any discrepancies between the manual and electronic measurements were used as a correction factor for the individual avatar representation of a participant. Within each block, participants were assessed in three domains of subjective experience and performance: unilateral goal-directed reaching, subjective perception of arm length, and subjective perception of embodiment. Each block started with a targeted reaching task, which comprised a gamified virtual reality experience, in which the participant fed animated animals (Fig. S1c). The animals were presented at four distances scaled to a participant’s extended arm length at 52%, 64%, 88%, and 100% extension. The four target animal distances were chosen so that the closer two were not affected by the GoGo manipulation, while the more distant two were located beyond the inflection point D and therefore within the range affected by the altered GoGo functionality.

Participants performed eight practice trials (two reaches to each distance) and 12 test trials (3 to each distance) in the first block; followed by 12 trials (3 to each distance) in the experimental block. In the practice trials, the target collider volume in which the food had to be released for the animal was visible as a yellow cube in front of an animal. This target volume was invisible in the actual trials. In baseline and experimental trials, each target position occurred three times in a randomized order, personalized to each participant.

After each start of the reaching game, a participant had a 5 second exploratory period to scan the surrounding environment and to look at their virtual hands before the first reaching trial commenced. When a reaching trial started, the grassy mound was bare. A blue readiness cube hovered in front of the participant (Fig S1c) and a food item (apple or carrot) appeared after a hand was moved into this volume. Keeping the hand inside the cube and closing the fingers by contacting the tips of the thumb and the index and middle fingers caused the food item to jump into the participant’s virtual hand. After pickup of the food item, an animal (randomised between a red panda, squirrel, or meerkat) appeared after a random delay (range 1s to 2s) at a specified target distance. Once the virtual hand was moved outside the readiness cube, the food item was attached to it and could not be released until the fingers were opened inside the animal’s target volume. A reaching trial ended with successful release and transfer of the food item. At this point, the animal was shown to nibble the food item and then disappear, with a new trial starting 1 second later.

The second part of a block aimed to evaluate a participant’s subjective reaching affordance – the perceived distance over which they could reach a target within a virtual environment. Instead of a grass mound, a long wooden table was presented in front of a participant: the tabletop was scaled in proportion to a participant’s height measurement. At the far end of the table, a stack of books and a tankard were placed as familiar-sized objects to help guide size and distance perception in the virtual scene. The hands and arms of a participant’s avatar were invisible, so the task was not influenced by current visual estimates of arm length. A red apple on a pedestal was presented at head level at variable locations on a participant’s straight-ahead midline (Fig. S1d). The anterior-posterior position of the apple was determined by a vector of 50 distance positions ranging from 0.5 to 3 times a participant’s arm lengths at logarithmically increasing spatial intervals. A participant’s task was to evaluate after each object presentation whether they were able to reach the apple with their real arm (without leaning forward). The apple remained visible until a verbal yes/no decision was reported and logged by the experimenter. The next apple was presented at a different distance. The distance of the first presentation was randomly determined, drawn from all 50 possible distances. All subsequent positions were determined by an adaptive staircase procedure (Parameter Estimation by Sequential Testing, PEST; Taylor & Creelman, 1967), which comprised 3 different step sizes depending on the trial number. In trials 1 to 7, after a ‘yes’ decision the next distance increased by 7 steps on the distance vector; after a ‘no’ decision it decreased by 7 steps. This step size was reduced to 3 following trials 8 to 12, and to 1 following trial 12. A maximum number of 30 trials was presented. Otherwise, the task was aborted when 5 yes/no reversals had occurred within the previous 7 trials, with counting towards this criterion starting after the 12^th^ trial. Staircase adjustments could not fall below the closest or go above the furthest distance. Another exceptional termination criterion was 5 repetitions of the highest or lowest possible distance. The subjective reaching affordance based on perceived physical arm length was determined as the average of the final three presentations.

### Questionnaire measures

The third part of a block consisted of the administration of five questions about the subjective experience during the reaching task of the first part. The questionnaire was read aloud by the experimenter, with the virtual reality headset was removed so that a participant was also able to read it at the same time. The questionnaire (Table 2) had one item on each of five themes: 1. general user experience, 2. perceived limb ownership, 3. perceived level of agency in terms of limb movement control, 4. any perceived tool-like properties of the limbs, and 5. participant’s suggestibility and answering bias. Each question was answered on a seven-point Likert scale (1: strongest disagreement, 4: neutral attitude, 7: strongest agreement). These items are based on existing best practice (Gonzalez-Franco & Peck, 2018) and have previously been used extensively with children in real and virtual bodily illusion studies (Cowie et al., 2013; Dewe et al., 2022; Filippetti & Crucianelli, 2019; Gottwald et al., 2021; Weijs et al., 2021).

**Table 2.** Embodiment questionnaire: questions and possible answers.

| Thematic category | Question | Disagreement | Neutral attitude | Agreement |
| --- | --- | --- | --- | --- |
| General user experience | While I was feeding the animals, I thought ... | ○ ... it was not so good (3)<br>○ ... I didn’t like it (2)<br>○ ... I hated it (1) | ○ ... it was neither good nor bad (4) | ○ ... I loved it (7)<br>○ ... I liked it (6)<br>○ ... it was ok (5) |
| Limb ownership | While I was feeding the animals, the virtual hands and arms felt like they were my own hands and arms, or belonged to me... | ○ ... not really (3)<br>○ ... not at all (2)<br>○ ... absolutely not (1) | ○ ... in between (4) | ○ ... lots and lots (7)<br>○ ... a lot (6)<br>○ ... a little bit (5) |
| Limb agency | While I was feeding the animals, I felt that I was in control of moving the virtual hands and arms ... | ○ ... not really (3)<br>○ ... not at all (2)<br>○ ... absolutely not (1) | ○ ... in between (4) | ○ ... lots and lots (7)<br>○ ... a lot (6)<br>○ ... a little bit (5) |
| Tool-like properties | While I was feeding the animals, the virtual hands and arms felt like a litter picker or grabber tool rather than part of my body ... | ○ ... not really (3)<br>○ ... not at all (2)<br>○ ... absolutely not (1) | ○ ... in between (4) | ○ ... lots and lots (7)<br>○ ... a lot (6)<br>○ ... a little bit (5) |
| Suggestibility and answering bias | While I was feeding the animals, my hair turned blue ... | ○ ... not really (3)<br>○ ... not at all (2)<br>○ ... absolutely not (1) | ○ ... in between (4) | ○ ... lots and lots (7)<br>○ ... a lot (6)<br>○ ... a little bit (5) |

### Movement data post-processing and reduction

Data logging of hand and head movements was performed by the inbuilt headset tracking facility, with a sampling rate of 62 - 71 Hz. The movement data acquired during the virtual reality reaching task were processed in MATLAB 2019b (Mathworks, Natick, MA) in a sequence of steps. Movement data logs were converted from text format into MATLAB binary format and segmented into the individual reaching trials based on event logs separating a trial into four phases: preparation, readiness, reaching, post-release. For data analysis, only the readiness and reaching phases were considered.

All movement trajectories were inspected and data gaps in movement trajectories due to intermittent loss of motion tracking were interpolated using a spline interpolation algorithm. Subsequently, all movement trajectories were smoothed using a dual-pass 4^th^ order Butterworth low-pass filter with a cutoff frequency of 6 Hz.

For the determination of motor performance in each reaching trial, several spatial and temporal parameters were extracted based on the position and velocity time series along the forward reaching (AP, anteroposterior) axis. In a custom semi-automatic algorithm, the boundaries of a pre-reach baseline period and a post-reach steady state were specified manually. Onset of a reaching movement was determined as the first frame outside the baseline period, which exceeded at threshold of four times the standard deviation of AP velocity during the baseline period.

The moment a target animal appeared served as a time reference for any temporal parameters (a negative time-to-event means that the event happened before the onset of the target animal).The position and timepoint of maximum AP arm extension (i.e. reaching distance) before an end-of-reach steady state was achieved - as well as the position, point in time, and magnitude of AP peak velocity - were determined automatically. Time to peak velocity was expressed as a proportion of the duration of maximum reach; the position of peak velocity was expressed as a proportion of maximum reach amplitude. Finally, as a measure of the smoothness of reaching movements we calculated spectral arc length as per (Balasubramanian et al., 2015) across the combined readiness baseline and reaching phases. Figure 1e shows as a function of time the position, velocity, and arm extension for an illustrative reaching trial of a single participant.

### Statistical Analysis

All statistical computations were performed in RStudio 2023.06.1+524 using R version 4.3.1. Ordinal scaled data such as the questionnaire ratings were aligned-rank-transformed (Wobbrock et al., 2011) before statistical analysis. We then used mixed repeated-measures Analyses of Variance (ANOVA) with between-subject factors age group (three levels: 5-to-7-year-olds, 8-to-10-year-olds, adults) and GoGo functionality (three levels: F-, F+, F++) and within-subject factor experimental block (two levels: baseline, GoGo). Target animal distance (four levels: 52%, 64%, 88%, 100% arm extension distance) was an additional within-subject factor for the movement-related performance parameters. Post-hoc single comparisons were performed following the Tukey procedure. Linear regressions were calculated and Spearman correlations between single parameters were performed using the parameters’ difference values between the baseline to the GoGo block.

### Data availability

All extracted parameter values for statistical analysis as well as supplementary materials are available from the figshare repository (https://doi.org/10.6084/m9.figshare.28130111).

## Supporting information

Supplemental results and demographics

## Funder acknowledgements

This research has been funded by ESRC ES/W003120/1 to DC, MG, SP. Thanks to all participating children, schools, and to the Life Science Centre, Newcastle.

