## Supplemental results and demographics for "Embodiment of a functionally altered virtual arm in adults and children"

Corresponding author:

Leif Johannsen

Cognitive and Experimental Psychology

### Methods

Table S1. Demographic information of all participants divided by age group, visual control gain group and gender. Averages (standard deviations) are reported. IOD: interocular distance; h/wk: hour per week; yrs: years; F-: 0.8 visual gain; F+: 1.2 visual gain; F++: 4.0 visual gain.

| Age group | Visual control gain group | Gender | N | Age (yrs) | Height (cm) | Arm length (cm) | IOD (cm) | Video game play (h/wk) |
| --- | --- | --- | --- | --- | --- | --- | --- | --- |
| 5-7 yrs | F- | Both | 25 | 7.2 (0.8) | 126.0 (7.6) | 38.0 (3.3) | 5.6 (0.8) | 7.5 (8.6) |
|  |  | Females | 12 | 7.0 (1.0) | 122.1 (7.6) | 36.8 (3.3) | 5.4 (0.9) | 4.4 (4.2) |
|  |  | Males | 13 | 7.3 (0.5) | 129.6 (5.7) | 39.1 (3.1) | 5.7 (0.7) | 10.4 (10.6) |
|  | F+ | Both | 24 | 7.0 (0.9) | 125.1 (8.2) | 37.8 (4.1) | 5.5 (0.6) | 7.4 (6.5) |
|  |  | Females | 20 | 6.9 (0.9) | 124.6 (8.8) | 37.5 (3.9) | 5.5 (0.5) | 7.8 (6.7) |
|  |  | Males | 4 | 7.1 (0.6) | 128.0 (3.6) | 39.0 (5.5) | 5.5 (1.1) | 5.8 (6.0) |
|  | F++ | Both | 21 | 7.0 (0.7) | 126.5 (6.6) | 37.5 (3.4) | 5.6 (0.7) | 6.7 (6.5) |
|  |  | Females | 10 | 6.8 (0.8) | 124.8 (6.8) | 36.3 (2.9) | 5.5 (0.8) | 4.4 (5.2) |
|  |  | Males | 11 | 7.2 (0.5) | 128.0 (6.4) | 38.5 (3.7) | 5.8 (0.6) | 8.8 (7.1) |
| 8 – 10 yrs | F- | Both | 30 | 9.5 (1.1) | 137.9 (7.3) | 42.7 (4.3) | 5.7 (0.7) | 9.2 (8.5) |
|  |  | Females | 15 | 9.6 (1.1) | 138.5 (7.6) | 43.0 (3.9) | 5.6 (0.6) | 8.3 (8.9) |
|  |  | Males | 15 | 9.5 (1.1) | 137.2 (7.2) | 42.5 (4.9) | 5.7 (0.8) | 10.2 (8.3) |
|  | F+ | Both | 35 | 9.0 (1.9) | 139.5 (9.6) | 42.6 (4.6) | 5.8 (0.7) | 8.9 (8.0) |
|  |  | Females | 15 | 9.4 (1.0) | 140.3 (10.4) | 43.7 (5.0) | 6.0 (0.3) | 8.5 (8.7) |
|  |  | Males | 20 | 8.7 (2.3) | 139.0 (9.2) | 41.9 (4.3) | 5.7 (1.0) | 9.2 (7.5) |
|  | F++ | Both | 36 | 9.2 (1.0) | 138.6 (8.9) | 42.6 (4.5) | 5.8 (0.7) | 10.0 (7.8) |
|  |  | Females | 14 | 9.2 (1.1) | 137.0 (9.2) | 42.1 (5.0) | 5.8 (0.8) | 7.2 (6.8) |
|  |  | Males | 22 | 9.3 (1.1) | 139.6 (8.8) | 42.8 (4.1) | 5.8 (0.7) | 11.8 (8.0) |
| >18 yrs | F- | Both | 31 | 22.3 (3.8) | 169.9 (8.2) | 49.7 (3.7) | 6.2 (0.4) | 4.1 (7.4) |
|  |  | Females | 24 | 21.8 (2.3) | 167.1 (4.5) | 48.5 (3.0) | 6.1 (0.3) | 2.5 (3.5) |
|  |  | Males | 7 | 24.2 (6.9) | 179.6 (11.0) | 53.9 (2.5) | 6.5 (0.5) | 9.6 (13.5) |
|  | F+ | Both | 27 | 22.4 (4.4) | 170.5 (8.6) | 50.2 (4.0) | 6.3 (0.5) | 4.8 (7.5) |
|  |  | Females | 15 | 21.2 (1.9) | 164.5 (3.4) | 48.3 (3.5) | 6.1 (0.4) | 3.1 (5.1) |
|  |  | Males | 12 | 23.7 (6.0) | 177.9 (7.1) | 52.6 (3.3) | 6.6 (0.6) | 7.0 (9.6) |
|  | F++ | Both | 30 | 22.6 (5.7) | 168.0 (8.0) | 49.4 (4.4) | 6.1 (0.6) | 5.4 (9.6) |
|  |  | Females | 24 | 22.1 (5.1) | 165.3 (5.2) | 48.3 (3.8) | 5.9 (0.5) | 6.3 (10.5) |
|  |  | Males | 6 | 24.3 (7.7) | 178.8 (8.1) | 53.8 (3.9) | 6.9 (0.4) | 1.8 (2.4) |

Figure S1. A.) a relationship between real (Rr) and virtual (Rv) hand position: an inflection point (D) at 2/3 arm length (A) was chosen as a comfortable distance for a transition between normal and altered functionality. As the real arm extended further than D (R𝑟 > D), the virtual arm length began to change in linear proportion to the real distance travelled. The maximum was a multiple, M, of the arm length: in the F- condition M=0.8; F+ M=1.2; F++ M=4.0. B.) Illustrations of the reaching sequence for a normal (I-IV) and a F++ (V-VIII) reach: moving into the ready volume triggers the appearance of a food item, grasping the food item triggers onset of a target animal, the forward reach is performed and the food item released, the animal nibbles the food item and disappears. C.) Illustration of the stimulus used in the reaching affordance task. An apple on a pedestal is presented in front of the participant, who then needs to reply whether the apple is in reach or not considering their physical arm length. A participant’s virtual hands and arms are not visible. D.) Anteroposterior position and velocity for an illustrative reaching trial of a single participant. Vertical lines indicate (from left to right) the movement preparation period and onset of the target animal, the start of the forward reach, peak velocity, and the end of forward reach.

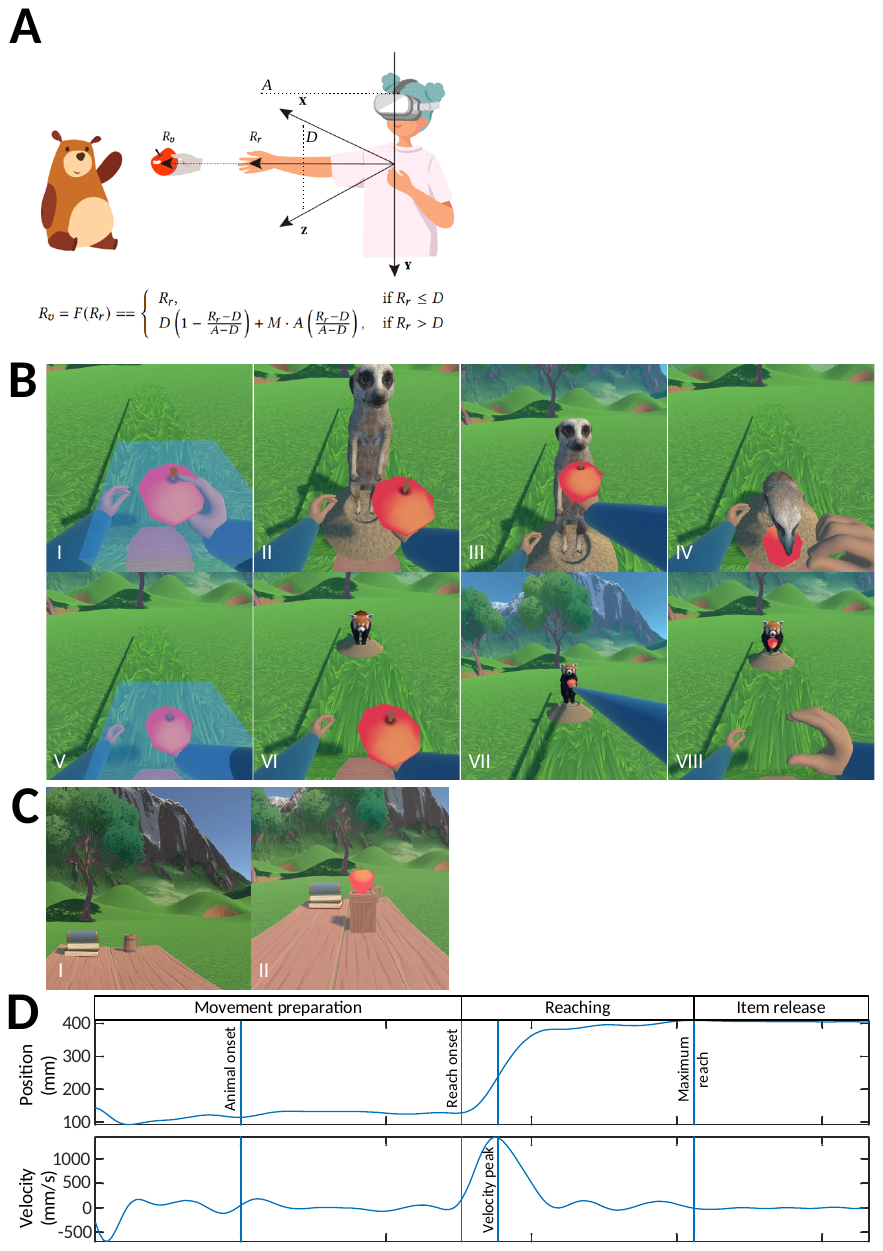

#### Calculation of surrogate visual target distance

The distance at which targets were visually presented was dependent on participants’ arm length and thus in absolute terms was greater for older (and taller) participants, who therefore reached for visually very distant targets in the F++ condition. While the virtual environmental coordinates at which targets were presented were not recorded, a surrogate measure was created based on participants’ physical arm length measurements and a body-scaling ratio.

First, target distance conditions in percentage arm length were multiplied by the measurement of a participant’s actual arm length to approximate the target locations in virtual space for each participant. Second, the farthest distance that was encountered by the participant with the longest arm would appear disproportionately more distant to the individual with the shortest arm length under the assumption that an individual’s visual distance perception might be based on a body-scaled frame of reference. To control for any body size-dependent perceptual biases, the approximated visual distances for each participant were normalized to the shortest arm length encountered in the study. For example, in the F++ condition the specific metric distance of four times arm length of an adult might resemble six times the arm length of the shortest child. Therefore, the approximated distances were multiplied by a factor resembling the arm length ratio of a participants’ arm length over the arm length of the shortest participant.

### Results

Table S1. Overview of significant effects and interactions by the subjective experience and embodiment performance parameters. Age: Age Group; Funct.: Functionality; Block: experimental block.

|  | Parameter | | | | | |
| --- | --- | --- | --- | --- | --- | --- |
|  | User  experience | Ownership | Agency | Tool  quality | Suggestion | Reaching  affordance |
| Age | **F(2,500)=4.82,**  **p=0.008,**  **pη^2^=0.02** | F(2, 500)=2.57,  p=0.08,  pη^2^=0.01 | F(2, 500)=0.98 ,  p=0.37,  pη^2^=0.004 | F(2, 500)=1.82,  p=0.16,  pη^2^=0.007 | **F(2, 500)=90.13,**  **p<0.001,**  **pη^2^= 0.26** | **F(2,249)=27.08,**  **p<0.001,**  **pη^2^=0.18** |
| Funct. | F(2,500)=0.77,  p=0.46,  pη^2^=0.003 | **F(2,500)=3.45,**  **p=0.03,**  **pη^2^=0.01** | F(2,500)=0.69,  p=0.50,  pη^2^=0.003 | **F(2,500)=5.70,**  **p=0.004,**  **pη^2^=0.02** | **F(2,500)= 4.33,**  **p=0.01,**  **pη^2^= 0.02** | F(2,)=0.08,  p=0.92,  pη^2^=0.0007 |
| Age * Funct | F(4, 500)=0.24,  p=0.92,  pη^2^=0.002 | **F(4,500)=4.09,**  **p=0.003,**  **pη^2^=0.03** | **F(4,500)=3.59,**  **p=0.007,**  **pη^2^=0.03** | F(4,500)=1.14,  p=0.33,  pη^2^=0.009 | **F(4,500)= 3.53,**  **p=0.007,**  **pη^2^=0.03** | F(4,)=0.45,  p=0.77,  pη^2^=0.007 |
| Block | F(1, 500)=0.05,  p=0.82,  pη^2^=0.0001 | F(1, 500)=1.97,  p=0.16,  pη^2^=0.004 | **F(1,500)=8.67,**  **p=0.003,**  **pη^2^=0.02** | F(1, 500)=2.21,  p=0.14,  pη^2^=0.004 | **F(1,500)=10.21,**  **p=0.001,**  **pη^2^=0.02** | **F(1,249)= 27.75,**  **p<0.001,**  **pη^2^=0.10** |
| Age * Block | F(2, 500)=0.35,  p=0.70,  pη^2^=0.001 | **F(2,500)=4.03,**  **p=0.02,**  **pη^2^=0.02** | F(2,500)=0.67,  p=0.51,  pη^2^=0.003 | F(2,500)=0.94,  p=0.39,  pη^2^=0.004 | **F(2,500)= 6.26,**  **p=0.002,**  **pη^2^=0.02** | F(2,)=2.15,  p=0.12,  pη^2^=0.02 |
| Funct. * Block | F(2, 500)=0.25,  p=0.78,  pη^2^=0.001 | **F(2,500)=4.46,**  **p=0.01,**  **pη^2^=0.02** | F(2, 500)=0.39,  p=0.68,  pη^2^=0.002 | F(2, 500)=1.84,  p=0.16,  pη^2^=0.007 | **F(2,500)=4.29,**  **p= 0.01,**  **pη^2^= 0.02** | **F(2,249)=6.42,**  **p=0.002,**  **pη^2^=0.05** |
| Age * Funct.  * Block | F(4,500)=0.13,  p=0.97,  pη^2^=0.001 | F(4,500)=1.12,  p=0.34,  pη^2^=0.009 | F(4,500)=0.55,  p=0.70,  pη^2^=0.004 | F(4,500)=2.06,  p=0.09,  pη^2^=0.02 | **F(4,500)=2.51,**  **p=0.04,**  **pη^2^= 0.02** | F(4,249)=0.63,  p=0.64,  pη^2^=0.01 |

### User experience in the game was rated slightly higher by younger children (mean=6.35, SD 0.89) and adults (mean=6.24, SD 0.89) than older children (mean=6.12, SD 0.78).

### Ownership was reported lower for the F++ arm, and especially so in adults, with slightly higher ratings in the baseline experimental block (mean=1.99, SD 1.59) compared to the GoGo block (mean=1.79, p=1.49; t(500)=3.20, p=0.002). The F- group (mean=5.17, SD 1.58) gave higher ratings than the F+ (mean=4.83, SD 1.63; t(500)=1.95, p=0.05) and F++ (mean=4.82, SD 1.61; t(500)=2.49, p=0.01) groups.

### Regarding the sense of agency, an interaction between age group and functionality group revealed that in younger children, the F+ group showed lower sense of agency than the F++ group (t(500)=2.27, p=0.02). In the group of adults, however, the F- group showed greater sense of agency compared to the F++ visual gain group (t(500)=3.76, p<0.001). No differences were observed for the older children. An interaction between age group, visual gain group, and experimental block was caused by the fact that the young children and the adults did not change their ratings after the GoGo experience, while the older children in the 4.0 visual gain group reduced their ratings (t(500)=2.05, p=0.04).

### Motor control

#### Reaching duration

An age effect revealed that reaching duration was shortest in the adults (mean=0.92 s, SD 0.24), followed by the older children (mean=1.07 s, SD 0.24; t(250)=4.35, p<0.001). Reaching duration lasted the longest in the younger children (mean=1.16 s, SD 0.24; t(250)=2.59, p=0.03). With respect to target distance, the 52% arm length distance resulted in the shortest reaches (mean=0.90 s, SD 0.29), followed by 64 % arm length (mean=0.95, SD 0.28; t(250)=2.58, p=0.05), by 88 % arm length (mean=1.14, SD 0.33; t(250)=9.64, p<0.001), and by 100% (mean=1.21 s, SD 0.34; t(250)=3.21, p=0.008). Figure S2 shows reaching duration for each age group as a function of functionality group, experimental block, and target distance.

Separate ANOVAs performed to resolve the age and functionality group interaction showed that only the functionality groups of the younger children differed against each other (F(2,67)=5.44, p=0.007, pη^2^=0.14). The F+ functionality group took longest for an average reach (mean=1.31s, SD 0.27) compared to the other two functionality groups (F-: mean=1.10s, SD 0.26, t(67)=2.79, p=0.02; F++: mean=1.08 s, SD 0.27, t(67)=2.90, p=0.01).

Examining the interaction between functionality group and target distance, we found that all functionality groups showed a significant effect of target distance (all F(3,255)>21.11, all p<0.001, all pη^2^>0.20). The F- functionality groups showed increases in movement duration between the first and second and the third target positions (both t(85)> 2.74, both p<0.04). The F+ functionality groups showed increases between the second, third, and fourth target positions (both t(85)> 3.17, both p<0.01). The F++ functionality groups showed an increase between the second and third target positions only (t(86)=5.92, p<0.001). All functionality groups had in common that the increase in distance between the target positions at 64% and 88% arm length increased reaching duration, but the functionality groups differed with respect to a difference between the first and second, and third and fourth target distances.

Fig. S2. Reaching duration as a function of age group, functionality group, block, and target distance condition. F-: reduced function; F+: increased function; F++: enhanced increased function. Error bars show the confidence interval. Asterisks indicate statistically significant single comparisons (p<0.05).

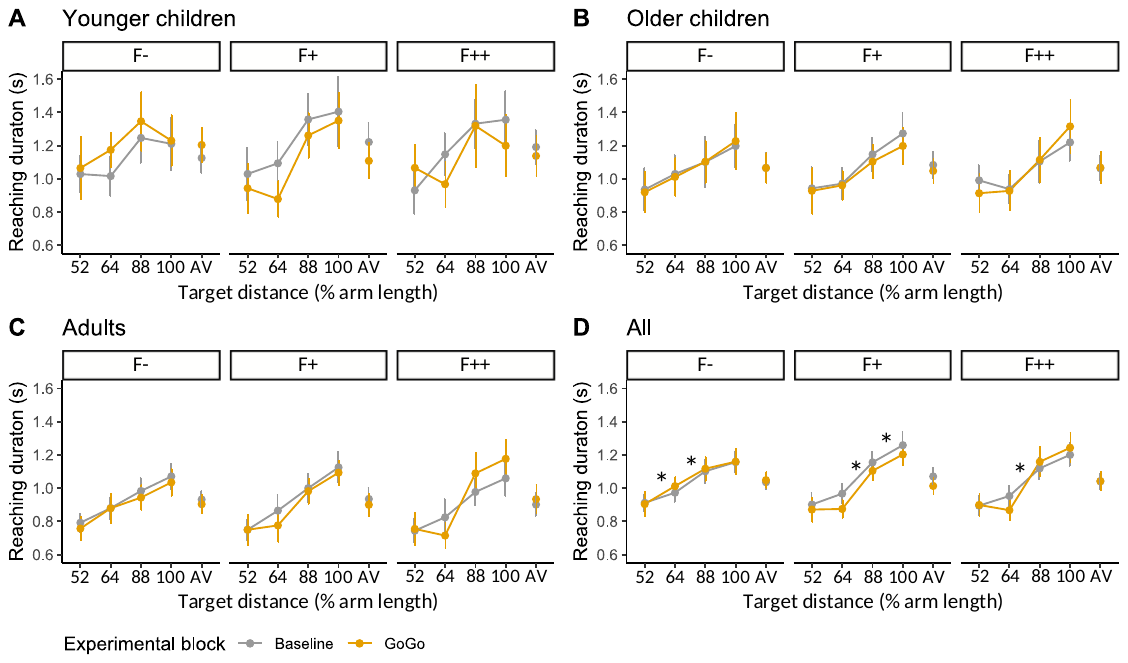

#### Peak velocity

Peak velocity increased with age, GoGo condition, and target distance. Adults (mean=586 mm/s, SD 132) were fastest, followed by older children (mean=545 mm/s, SD 131) and then younger children (mean=463 mm/s, SD 132). Peak velocity was higher in the GoGo block (mean=542 mm/s, SD 156) than at baseline (mean=520 mm/s, SD 143). Across the four target distances linear trends of peak velocity were observed for younger children (t(67)=5.79, p<0.001), for older children (t(98)=10.85, p<0.001), and for adults (t(85)=22.08, p<0.001).

To resolve the two 2-way interactions involving age group, post-hoc comparisons between age groups were calculated. Only the older children showed an increase in peak velocity in the experimental block with modified functionality (t(250)=3.23, p=0.02), while the younger children and the adults did not alter peak velocity between the two blocks (both t(250)<2.58, both p>0.11). Regarding the effect of target distance, the younger children increased peak velocity between the first and second target distances (t(67)=3.43, p=0.006; second vs third: t(67)=1.49, p= 0.45; third vs fourth: t(67)= 2.58, p=0.06) only. The older children showed increases in peak velocity between the first and second target position (t(98)=3.80, p=0.001) and the second and third (t(98)=6.92, p=<.001), but not between the third and fourth (t(98)=0.035, p=1.00). The adults, on the other hand, showed increases in peak velocity between all distances (all t(85)=5.11, all p<0.001).

Separate ANOVAs and post-hoc comparisons were performed for each functionality group to resolve the 3-way interaction between functionality group, experimental block, and target animal distance. The ANOVAs are reported in the article’s main text. Post-hoc comparisons for each functionality condition indicated that for the F- functionality groups the magnitude of peak velocity increased only from the closest to the second closest target position and from the second target position on remained on the same level for the two distances affected by the functionality modification (t(83)=3.17, p=0.01). In the F+ functionality groups a rise in the magnitude of peak velocity was present from the first to the second target position (t(83)=4.95, p<0.001) and from the second to the third (t(83)=4.03, p<0.001) but remained on the same level from position 3 to 4 (t(83)=1.46, p=0.47). A similar pattern was observed for the F++ functionality groups (position 1 to 2: t(84)=4.47, p<0.001; position 2 to 3: t(84)=8.99, p<0.001; position 3 to 4: t(84)=1.16, p=0.65). However, the effect size of the increase in peak velocity from target position 2 to 3 was greater in the F++ (F(2,84)=80.80, p<0.001, pη^2^=0.49) compared to the F+ functionality groups (F(2,83)=16.28, p<0.001, pη^2^=0.16).

Interestingly, the F+ groups in contrast demonstrated an effect of block irrespective of target distances (F(1,83)=6.04, p=0.02, pη^2^= 0.07). A main effect of block was not observed for the F- (F(1,83)=1.25, p=0.27, pη^2^=0.01) and F++ groups (F(1,84)=1.47, p=0.23, pη^2^=0.02). As an alternative approach to investigate the 3-way interaction, additional ANOVAs for each experimental block showed no effects or interactions involving functionality groups at baseline (both F<2.31, both p>0.10), but an interaction between functionality group and target distance in the GoGo block (F(6,750)= 17.47, p<0.001, pη^2^=0.12).

#### Absolute time to peak velocity (aTPV)

Faster reaches were observed in adults, in the F++ and F- groups, and for near targets. The adults showed the shortest aTPV (mean=0.33 s, SD 0.14), followed by the older children (mean=0.43 s, SD 0.14; t(250)=4.95, p<0.001) and the younger children (mean=0.49 s, SD 0.14; t(250)= 2.93, p= 0.01). The shortest aTPV occurred in the F++ group (mean=0.38, SD 0.14), followed by the F- group (mean=0.43, SD 0.14; t(250)= 2.21, p=0.07), and the F+ group (mean=0.45, SD 0.14; t(250)=1.22, p=0.44). The closest target distance showed the shortest aTPV (mean=0.38 s, SD 0.19), followed by the second closest target position (mean=0.39, SD 0.19; t(250)=0.46, p=0.97), the forth (mean=0.45, SD 0.21; t(250)=3.72, p=0.001) and the third target positions (mean=0.46, SD 0.23; t(250)=0.81, p=0.85).

Younger children changed aTPV substantially with target position and Gogo condition, while adults held it remarkably constant. This is seen in the interaction between age group, functionality group and block (Fig. S3). Peak velocity occurred later for further targets in young children (F(3,201)=10.28, p<0.001, pη^2^=0.13), but not older children (F(3,294)=2.50, p=0.06, pη^2^=0.02) or adults (F(3,255)= 1.51, p=0.21, pη^2^=0.02). Further, the younger children showed an interaction between functionality group and block (F(2,67)=7.47, p=0.001, pη^2^=0.18): aTPV in the Gogo block increased for the F- group (F(1,24)=8.52, p= 0.008, pη^2^=0.26), but tended to reduce for the F+ (F(1,23)=4.40, p=0.05, pη^2^=0.16) and F++ (F(1,20)=2.42, p=0.14, pη^2^=0.11) groups. Older children showed a similar, but weaker pattern: an interaction between functionality and block (F(2,98)=5.28, p=0.007, pη^2^=0.10)) with reduced aTPV in the F++ group Gogo block (F(1,35)=7.61, p=0.009, pη^2^=0.18). No effect of block was found for the F- (F(1,29)=2.95, p=0.10, pη^2^=0.09) and F+ (F(1,34)=0.13, p=0.72, pη^2^=3.81e-03) groups. Adults, in contrast, showed no interaction between functionality group and block (F(2,85)= 0.38, p=0.68, pη^2^=8.94e-03). These changes to aTPV reveal a tendency for children to spend longer in acceleration for the subfunctional F- arm. Irrespective of age group, an effect of experimental block was not observed in any of the three functionality groups (all F<1.08, all p>0.31, all pη^2^<0.03).

Fig. S3. Absolute time to peak velocity as a function of age group, functionality group, block, and target distance condition. F-: reduced function; F+: increased function; F++: strongly increased function. Error bars show the confidence interval. Asterisks indicate statistically significant single comparisons (p<0.05).

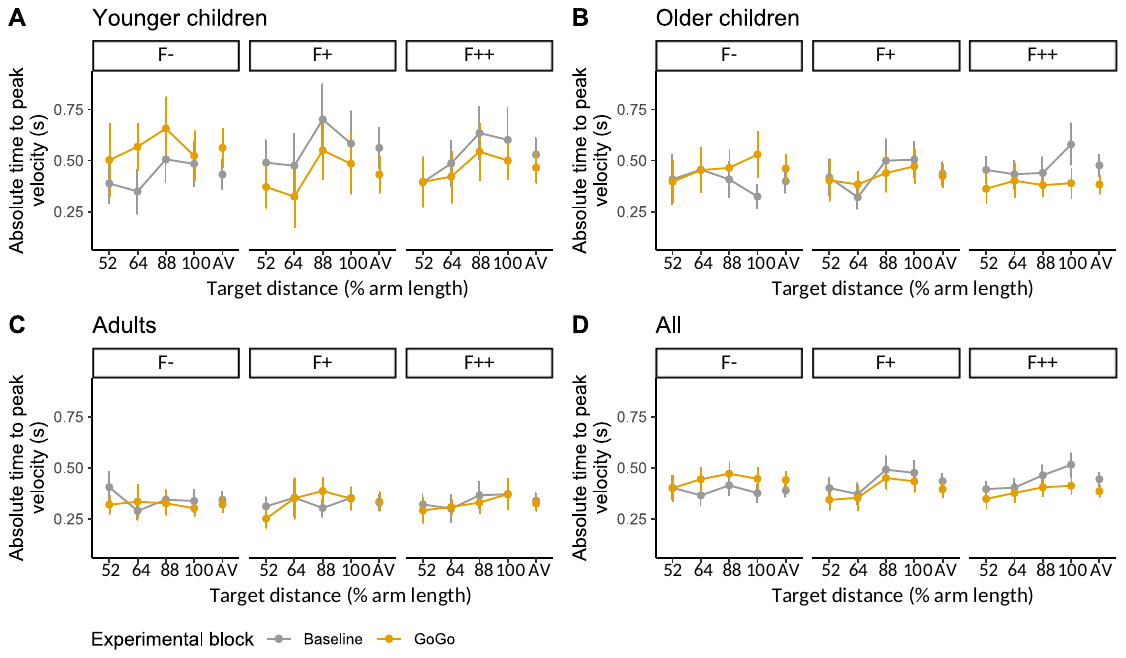

#### Relative time to peak velocity (rTPV)

An interaction between block and target distance was caused by the closest target distance showing later rTPV than the other three target distances in the baseline block (all t(250)>3.11 , all p<0.04). In the GoGo block, no differences between target positions occurred. Both younger (F(2,67)=3.13, p=0.05, pη^2^= 0.09) and older children (F(2,98)= 5.75, p=0.004, pη^2^= 0.11) showed an interaction between functionality group and experimental block, while adults did not (F(2,85)=0.5851, p=0.56, pη^2^= 0.01).

The relative time to peak velocity as a proportion of the reach duration was shortest in the adults (mean=37.7%, SD 8.03) compared to the younger (mean=42.1%, SD 8.03) and older children (mean=41.3%, SD 8.03). The shortest rTPV (mean=37.6%, SD 12.8) occurred at the greatest target distance compared to the third (mean=40.1%, SD 13.3), second (mean=40.8%, SD 13.6), and the closest target distances (mean=42.8%, SD 14.3). Breaking down further, younger children in the F- group GoGo block (where forward movement was impeded) showed a tendency for a longer accelerative phase (increased rTPV; F(1,24)=3.08, p=0.09, pη^2^= 0.11). Older children also showed this (F(1,29)=4.72, p=0.04, pη^2^= 0.14), along with a reduction in the F++ group (F(1,35)=6.48, p=0.02, pη^2^= 0.16). In summary, adults were very consistent in their control of the timing of reaching by keeping the absolute timepoint of peak velocity constant irrespective of the target distance and the functionality condition, which resulted in a reduction of rTPV as reaching duration increased with target distance. Both groups of children responded with a complementary strategy by keeping rTPV constant, which means they timed peak velocity at a later point for more distant targets with longer reach duration. Nevertheless, differences between the two groups of children occurred, with an influence of target distance on timing in the older children but no influence in the younger children.

#### Absolute distance to peak velocity (aDPV)

The absolute distance to peak velocity was greater in the older children (mean=250, SD 42) compared to the younger children (mean=230, SD 42; t(250)=3.19, p=0.005) and the adults (mean=234, SD 42; t(250)=2.74, p=0.02). Between the functionality groups, the F++ groups tended to show the shortest distance to peak velocity (mean=228, SD 43) compared to the F- (mean=243, SD 42; t(250)=2.26, p=0.06) and F+ groups (mean=242, SD 42; t(250)=2.11, p=0.09). Distance to peak velocity increased from each target position to the next (52%: mean=209, SD 45; 64%: mean=222, SD 46; 88%: mean=256, SD 49; 100%: mean=264., SD 45; all t’s>3.82, all p’s<0.001).

Separate ANOVAs were performed to resolve a 4-way interaction between age group, functionality group, experimental block and target distance (Fig. S4). All three age groups showed an interaction between functionality group, experimental block and target distance (all F>3.09, all p<0.006, all pη^2^>0.06). In the baseline block, the younger children (F(3, 201)=24.31, p<0.001, pη^2^=0.27) and the adults (F(3,255)=110.95, p<0.001, pη^2^=0.57) demonstrated an effect of target distance only. In contrast, the older children showed an effect of target distance (F(3,294)=44.67, p<0.001, pη^2^=0.31) and an interaction between functionality group and target distance (F(6,294)=3.76, p=0.001, pη^2^=0.07). Post-hoc comparisons showed that in the 0.8 functionality group of the older children, the increments between distances were not significant (all t(29)<2.19, all p>0.15). For the F+ functionality group, the increments from the first to the second (t(34)=2.88, p=0.03) and to the third (t(34)=3.90, p=0.002) were significant, while the third and the fourth were not different (t(34)=1.15, p=0.66). In the F++ functionality group of the older children, the third and fourth target distances (t(35)=5.11, p<0.001) differed only.

In the Gogo block, the younger children showed an effect of target distance (F(3,201)=48.0, p<0.001, pη^2^=0.42) and an interaction between functionality group and target distance (F(6,201)=8.16, p<0.001, pη^2^=0.20). While the F- functionality group of the younger children showed no effect of target distance (F(3,72)=2.48, p=0.07, pη^2^= 0.09), the F+ functionality group, on the other hand, did show an effect of target distance (F(3,69)=15.30, p<0.001, pη^2^=0.40).Post-hoc comparisons expressed that an increase in peak velocity distance occurred between the second and third target positions (t(23)=4.339, p=0.001). The F++ functionality group also demonstrated an effect of target distance (F(3,60)=50.39, p<0.001, pη^2^=0.72). An increase in peak velocity position occurred again between the second and third target position (t(20)=7.16, p<0.001).

The older children showed an effect of functionality group (F(2, 98)=3.46, p=0.04, pη^2^= 0.07), an effect of target distance (F(3, 294)= 79.67, p<0.001, pη^2^= 0.45), and an interaction between the two factors (F(6,294)=5.40, p<0.001, pη^2^=0.10). The F- functionality group of the older children showed an effect of target distance (F(3, 87)=10.72, p<0.001, pη^2^= 0.27). An increase in the peak velocity distance occurred between the first and second target positions (t(29)=2.99, p=0.03). Also the F+ functionality group was influenced by target distance (F(3,102)=46.54, p<0.001, pη^2^= 0.58). An increase in peak velocity distance occurred between the second and third target positions (t(34)=6.44 p<0.001). In the same manner, the F++ functionality group showed an effect of target distance (F(3,105)=34.38, p<0.001, pη^2^=0.50). An increase in peak velocity position occurred again between the second and third target position (t(35)=6.72, p<0.001).

Like the older children, did the adults show an effect of functionality group (F(2,85)=4.81, p=0.01, pη^2^=0.10), an effect of target distance (F(3, 255)=150.31, p<0.001, pη^2^=0.64), and an interaction between the two factors (F(6,255)=13.15, p<0.001, pη^2^=0.24). The adult F- functionality group showed an effect of target distance (F(3,90)=14.05, p<0.001, pη^2^=0.32) caused by an increase between the first and second target position (t(30)=2.94, p=0.03). Similarly behaved the F+ functionality group (F(3,78)= 66.37, p<0.001, pη^2^=0.72) caused by increases from the first to the second (t(26)=4.18, p=0.002) and from the second to the third target positions (t(26)=9.48, p<0.001). Finally, the F++ functionality group demonstrated an effect of target position (F(3,87)=84.95, p<0.001, pη^2^=0.75). An increase in peak velocity position was apparent between the second and third target positions (t(29)=9.99, p<0.001) only.

An alternative approach for resolving the 4-way interaction between age group, functionality group, experimental block and target distance was also attempted. In the groups of younger children, the 0.8 visual gain group showed an effect of target distance only but neither an effect of experimental block nor an interaction between experimental block and target distance. No differences existed between the two experimental blocks for any of the four target distances (all t(24)<2.43, all p>0.27). In contrast, the F+ and F++ visual gain groups both demonstrated interactions between experimental block and target distance (both F>2.89, both p<0.04, both pη^2^>0.11). The F+ visual gain group demonstrated an increase in peak velocity distance for the third target position (t(23)=3.79, p=0.02), while the F++ visual gain group showed a reduction in the distance at the first target position (t(20)=3.36, p=0.05).

In the groups of older children, the F- visual gain group showed an effect of target distance only and no differences between experimental blocks for the four target positions (all t(29)<2.18, all p>0.39). The F+ visual gain group showed effects of experimental block and target distances, but no interaction between the two factors. Compared to the baseline block (mean=239, SD 51) was the peak velocity amplitude more distant in the second experimental block with modified visual gain (mean=259, SD 34; F(1,34)=7.61, p=0.009, pη^2^=0.18). The F++ visual gain group showed an effect of target distance and an interaction between experimental block and target distance (F(3,60)=9.16, p<0.001, pη^2^=0.31). Post-hoc comparisons, however, showed no differences between the experimental blocks for any target distance (all t(35)<2.47, all p>0.24).

In the groups of adults, the F- visual gain group also demonstrated an effect of target distance and an interaction between experimental block and target distance but no significant post-hoc comparisons between experimental blocks for each target distance were found (all t(30)<1.06, all p>0.96). The F+ visual gain group also showed an effect target distances and an interaction between experimental bock and target distance (F(3,78)=6.53, p<0.001, pη^2^=0.20). No significant post-hoc comparisons between experimental blocks for each target distance were found (all t(30)<1.65, all p>0.71). Lastly, also for the F++ visual gain group an effect of target distance and an interaction between experimental block and target distances were found (F(3,60)=9.16, p<0.001, pη^2^=0.31). Post-hoc comparisons between the experimental blocks for each target position were not significant (all t(29)>2.72, all p>0.16).

Fig. S4. Absolute distance to peak velocity as a function of age group, functionality group, block, and target distance condition. F-: reduced function; F+: increased function; F++: enhanced increased function. Error bars show the confidence interval. Asterisks indicate statistically significant single comparisons (p<0.05).

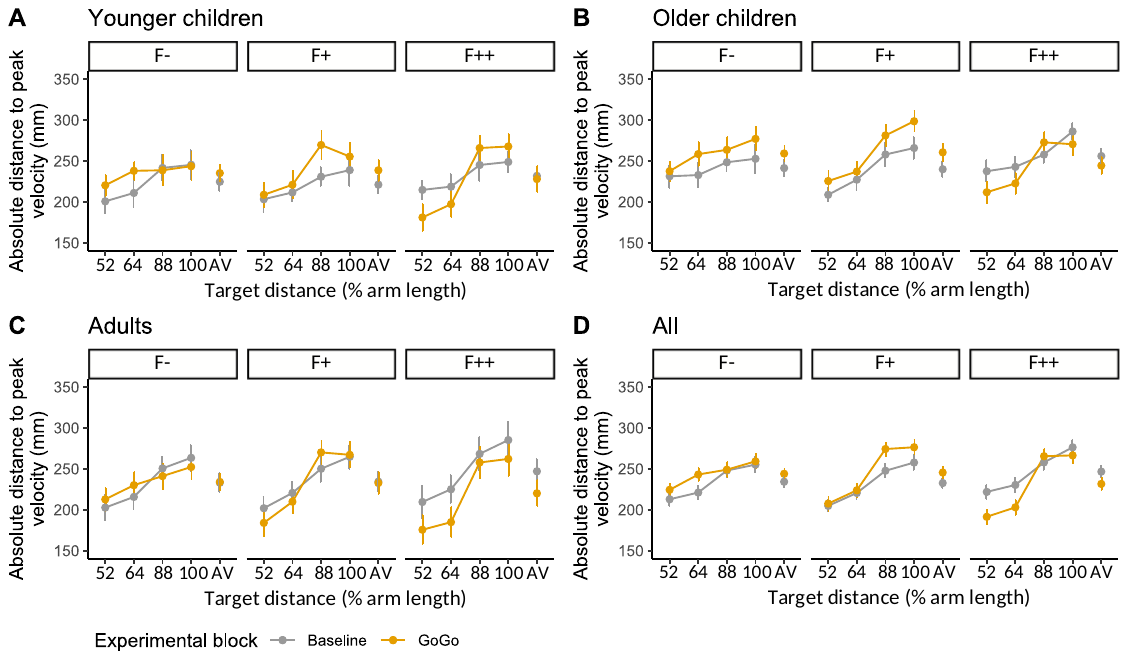

#### Relative distance to peak velocity (rDPV)

The relative distance to peak velocity increased with target distance (52% target distance mean=77.1 %, SD 8.47; 64% TD mean=74.4 %, SD 7.63; 88% TD mean=71.0 %, SD 7.16; 100% TD mean=68.9 %, SD 7.04). Adults (mean=68.0%, SD 5.76) reached peak velocity proportionally closer than younger children (mean=75.6 %, SD 5.77; t(250)= 8.15, p<.0001) and older children (mean=75.0 %, SD 5.77; t(250)= 8.27, p<0.0001). In the older children, we observed an effect of target distance (F(3,294)= 31.90, p<0.001, pη^2^=0.25), and an interaction between functionality group and block (F(2,98)= 5.95,p=0.004, pη^2^=0.11). Adults showed even stronger scaling through an effect of target distance (F(3,255)=106.07, p<0.001, pη^2^=0.56) and an interaction between functionality group, block and target distance (F(6,255)= 3.90, p<0.001, pη^2^=0.08).

In the F- group of the older children, a decreasing linear trend across the four target distances at baseline (t(29)=4.95, p<0.001) was not maintained in the GoGo block (t(29)=1.33, p=0.20) , while the F+ and F++ groups showed a linear decreasing trend across the four target distances in both blocks (F+ baseline: t(34)=4.21, p<0.001; F+ GoGo: t(34)=3.41, p=0.002; F++ baseline: t(35)=2.95, p=0.006; F++ GoGo: t(35)=4.71, p<0.001).

#### Smoothness

Smoothness was affected by age group, with the adults showing smoothest reaching (mean=-1.84, SD 1.30), followed by older (mean=-3.04, SD 1.31), and younger children (mean=-4.26, SD 1.31): all three groups differed from each other (all t>5.98, all p<0.001). Smoothness was affected by an interaction of functionality group, block, and target distance. At baseline, only an effect of target distance (F(3,768)=4.55, p=0.004, pη^2^= 0.02) was observed (an upwards linear trend with increasing target distance, t(256)=2.46, p=0.01). The GoGo block also showed an effect of functionality group (F(2,256)=10.17, p<0.001, pη^2^= 0.07) and an interaction between functionality and target distance (F(6, 768)=6.90, p<0.001, pη^2^= 0.05). The effect of target distance was only present in the increased function groups (F+: F(3,255)=7.62, p<0.001, pη^2^=0.08; F++: F(3,258)=14.18, p<0.001, pη^2^= 0.14) as linearly increasing trends (F+: t(85)=3.73, p<0.001; F++: t(86)=5.26, p<0.0001) with reduced smoothness in the two closer target positions likely reflecting some corrections for over-reaches when encountering enhanced arm length. This could be due to overgeneralisations in the motor control strategy based on the functionality context.

### Reaching affordance

The mean number of trials until a threshold was reached was 16.2 trials (range=6 to 30 trials). The maximum number of trials criterion where the staircase procedure was aborted was reached in one participant. Figure S5 shows the distributions of reaching affordance for each age group and functionality group.

Compared to the adults (mean=1.78, SD 0.39), the two groups of children (both t(249)>6.03, both p<0.001; young: mean=2.19, SD 0.39; older: mean=2.12, SD 0.39) overestimated the distances they were able to reach.
